# Downregulation of Satb1 is required to prevent autoimmunity by maintaining Tfh homeostasis

**DOI:** 10.64898/2026.02.03.703482

**Authors:** Mehrnoush Hadaddzadeh Shakiba, Maren Köhne, Tarek Elmzzahi, Luisa Bach, Doaa Hamada, Leonie Heyden, Anna Lindemann, Jannis B Spintge, Aleksej Frolov, Yuanfang Li, Lisa Holsten, Marlene Gottschalk, Celia Kho, Darya Malko, Collins Osei-Sarpong, Rebekka Scholz, Xiaoxiao Cheng, Anna Neubauer, Jonas Schulte-Schrepping, Yinshui Chang, Lorenzo Bonaguro, Heike Weighardt, Thomas Ulas, Joachim L Schultze, F. Thomas Wunderlich, Elena De Domenico, Axel Kallies, Zeinab Abdullah, Dirk Baumjohann, Marc D. Beyer

## Abstract

T follicular helper (Tfh) cells are a specialized subset of CD4⁺ T cells that localize to germinal centers (GC), where they provide critical help to B cells through the delivery of IL-21 and other cytokines. Here, we demonstrate that the tight control of the chromatin remodeler Special AT-rich sequence-binding protein 1 (Satb1) is key for this process, as overexpression of Satb1 drives lymphoproliferation and expansion of the T cell and B cell compartments in secondary lymphoid organs. Specifically, Satb1 overexpression induces a pronounced shift towards Tfh cell differentiation and increased GC formation accompanied by an increase in non-classed switched GC B cells and auto-antibody secretion. These findings highlight the importance of the precise regulation of Satb1 in fine-tuning CD4⁺ T cells and B cells responses and suggest a potential role for dysregulation of Satb1 in the pathogenesis of autoimmune disease such as systemic lupus erythematodes (SLE).

## INTRODUCTION

T follicular helper cells (Tfh) are a specialized subset of CD4^+^ T cells initially discovered in humans ^1^. Tfh cells play a crucial role in assisting B cells as they are essential for the formation of germinal centers (GCs), affinity maturation, and the development of high-affinity antibodies and memory B cells ^2, 3^. Thus, Tfh cells have emerged as key players in the regulation of adaptive immunity. Tfh cells are characterized by the expression of the transcription factor B cell lymphoma 6 (Bcl-6) and several cell surface markers, including C-X-C motif chemokine receptor type 5 (CXCR5), programmed cell death protein 1 (PD-1), and inducible co-stimulator (ICOS). These cells secrete interleukin-21 (IL-21), which plays a crucial role in promoting B cell differentiation and affinity maturation ^4, 5^. Previous studies have demonstrated that the upregulation of CXCR5, along with the downregulation of C–C chemokine receptor 7 (CCR7), is essential for the effective positioning of Tfh cells within the follicle and efficient B-cell help ^6^. The dysregulation of Tfh-B cell interaction can lead to abnormal B cell differentiation, potentially contributing to the development of autoimmune diseases such as systemic lupus erythematosus (SLE), rheumatoid arthritis (RA), and primary Sjögren’s syndrome (pSS) ^7, 8^. To regulate the GC response, T follicular regulatory (Tfr) cells—a specialized subset of regulatory T (Treg) cells—play a crucial role ^9^. They modulate the activity of Tfh cells and GC B cells, thereby influencing antibody production. Tfr cells express Forkhead box p3 (FOXP3), the transcription factor characteristic of Treg cells, while also sharing markers typical of Tfh cells, such as Bcl-6, CXCR5, ICOS, C-X-C chemokine ligand 13 (CXCL13), and PD-1 ^7, 8^.

It has been reported that loss of Tfh cell function results in impaired GC formation, defective class-switch recombination, and diminished production of high-affinity antibodies, thereby impairing the generation of long-lived plasma cells and memory B cells and contributing to pathological processes such as autoantibody production and tissue injury ^7^. Tfh cells, while specialized in supporting B cell responses, exhibit a remarkable degree of plasticity. Emerging evidence reveals the plastic nature of Tfh cells, demonstrating their capacity to derive from or transition into diverse helper T-cell subsets, including Th17 and Th2 cells, depending on environmental cues and the specific disease context ^10, 11^. This inherent flexibility underscores the need for robust regulatory mechanisms to stabilize the Tfh lineage. Precise regulation of Tfh cell differentiation, integrity, and localization is governed by intricate positive and negative feedback loops. At the core of this regulatory network is the transcription factor Bcl-6, which acts as a transcriptional repressor. Bcl-6 orchestrates Tfh differentiation through two key mechanisms: direct repression of alternative helper T-cell lineage drivers (T-bet, GATA-3, RORγt, STAT5, and Blimp1) and a repression-of-repressor mechanism that controls CXCR5 expression by releasing Klf2-mediated suppression of PD-1 and S1PR1 ^12^.

Recently, Chaurio et al. showed that repression or deletion of special AT-rich sequence-binding protein 1 (Satb1) in CD4^+^ T cells can enhance Tfh cell differentiation, promoting the formation of intratumoral tertiary lymphoid structures (TLS) and leading to reduced tumor growth in mice ^13^. Satb1 functions as a chromatin organizer and transcription factor, regulating gene expression by anchoring genomic regions to the nuclear matrix and recruiting transcription factors and chromatin modifying enzymes to target gene loci ^14, 15^. It plays a crucial role in lymphoid lineage specification of hematopoietic stem cells and thymocyte development, as well as in the differentiation of mature CD4^+^ T cell subsets and early B cell differentiation ^16^. Studies using Satb1-null mice revealed that thymocyte development failed mainly at the DP stage in the absence of Satb1 ^17^. In fact Satb1 has an essential role in positive and negative selection of thymocytes, and for the establishment of immune tolerance as its presence is necessary to shape the epigenetic landscape to enable Treg cell differentiation^15^. Satb1 plays a crucial role in directing the differentiation of CD4^+^ T cells, specifically by favoring the skewing of naïve CD4^+^ T cells into Th2 and Th17 cells, while simultaneously suppressing the development of Th1 and Treg cells ^18^. These findings support the hypothesis that the fine-tuned regulation of Satb1 expression and the downstream epigenetic events in CD4^+^ T cells are essentially controlling which types of T cells become dominant during a CD4^+^ T cell response ^18^. Recent studies have established Satb1 as a pivotal regulator of CD8⁺ T-cell fate, demonstrating its necessity for maintaining stem-like precursors and mitigating excessive effector differentiation, particularly in chronic infections and cancers ^19, 20^. Specifically, Heyden *et al*. showed that Satb1 is selectively enriched in precursor-exhausted CD8⁺ T cells and its downregulated is a prerequisite for their transition into terminal effector cells, revealing a dosage-dependent mechanism by which Satb1 restrains effector differentiation and modulates antiviral and antitumor immunity ^19^. However, its precise role in Tfh remains to be fully elucidated. Therefore, the main aim of this study is to characterize the functional impact of sustained expression of Satb1 in CD4^+^ T cells.

In this study, we observed lymphoproliferation, leading to lymphadenopathy, hepatomegaly, and splenomegaly in aged mice with sustained Satb1 expression in CD4^+^ T cells caused by a substantial expansion of the T cell compartment as well as B cells and myeloid cells. Overexpression of Satb1 increased the differentiation of CD4^+^ T cells towards a Tfh phenotype both *in vivo* and *in vitro* and resulted in an increase in GC formation. Characterization of the B cell population revealed that overexpression of Satb1 resulted in an accumulation of GC B cells but impaired the development of antigen-specific, class-switched B cells. Overall, our findings suggest that the precise regulation of Satb1 expression is a critical aspect of CD4^+^ T cell differentiation to fine tune the development of fully functional Tfh cells.

## RESULTS

### Sustained high levels of Satb1 during CD4^+^ T cell differentiation cause lymphoproliferation and an expansion of CD4^+^ T cells in aged Satb1-KI mice

Our previous work has indicated that Satb1 overexpression in CD4^+^ T cells can promote Th17 cell differentiation. This effect was characterized by an elevated proportion of IL-17⁺ and RORγt⁺ CD4^+^ T cells in the gut^18^, consistent with established mechanisms of Th17 cell development ^21^. To further investigate the influence of Satb1 overexpression in CD4⁺ T cells, we utilized the established Rosa26-STOP-Satb1 conditional model (in the following referred to as knockin (KI) mice) (**Supplementary Fig. 1a**) ^18^, which enables Cre-dependent Satb1 expression upon crossing to CD4-Cre mice. Additionally, we introduced the Foxp3-RFP reporter allele, allowing identification of Foxp3-expressing cells via RFP fluorescence.

To assess the influence of Satb1 overexpression on T cell dynamics, we quantified T cell populations in wild-type (CD4-Cre^wt/wt^, Rosa26-STOP-Satb1^tg/wt^, WT) and conditional CD4-Cre^tg/wt^, Rosa26-STOP-Satb1^tg/wt^ (Satb1 KI) mice across the lifespan of the animals. We observed a trend towards increased absolute T cell numbers in KI mice from 24 weeks on, with significant increases evident from week 35 on in the spleen and week 48 on in the mesenteric lymph nodes (mLN) (**Fig. 1a, b** and **Supplementary Fig. 1b**). Concomitantly, we observed a massive lymphoproliferation in aged Satb1 KI mice, resulting in demonstrable enlargement of the spleen and lymph nodes, including the mLNs and peripheral lymph nodes (pLNs) (**Fig. 1c**). This lymphoproliferative response was pronounced in both aged female and male KI mice. Hematoxylin and eosin (H & E) staining of spleen sections from Satb1 KI mice revealed significantly enlarged germinal center structures compared to WT mice (**Supplementary Fig. 1c**). In addition, Satb1 KI mice exhibited increased splenic weight relative to WT controls (**Supplementary Fig. 1d**). Furthermore, Satb1 KI mice exhibited a markedly reduced survival rate, with only 50% surviving to 52 weeks of age (**Fig. 1d**). These data support that a sustained expression of Satb1 in CD4^+^ T cells is affecting the size of secondary lymphoid organs and ultimately results in reduced lifespan.

**Figure 1:**
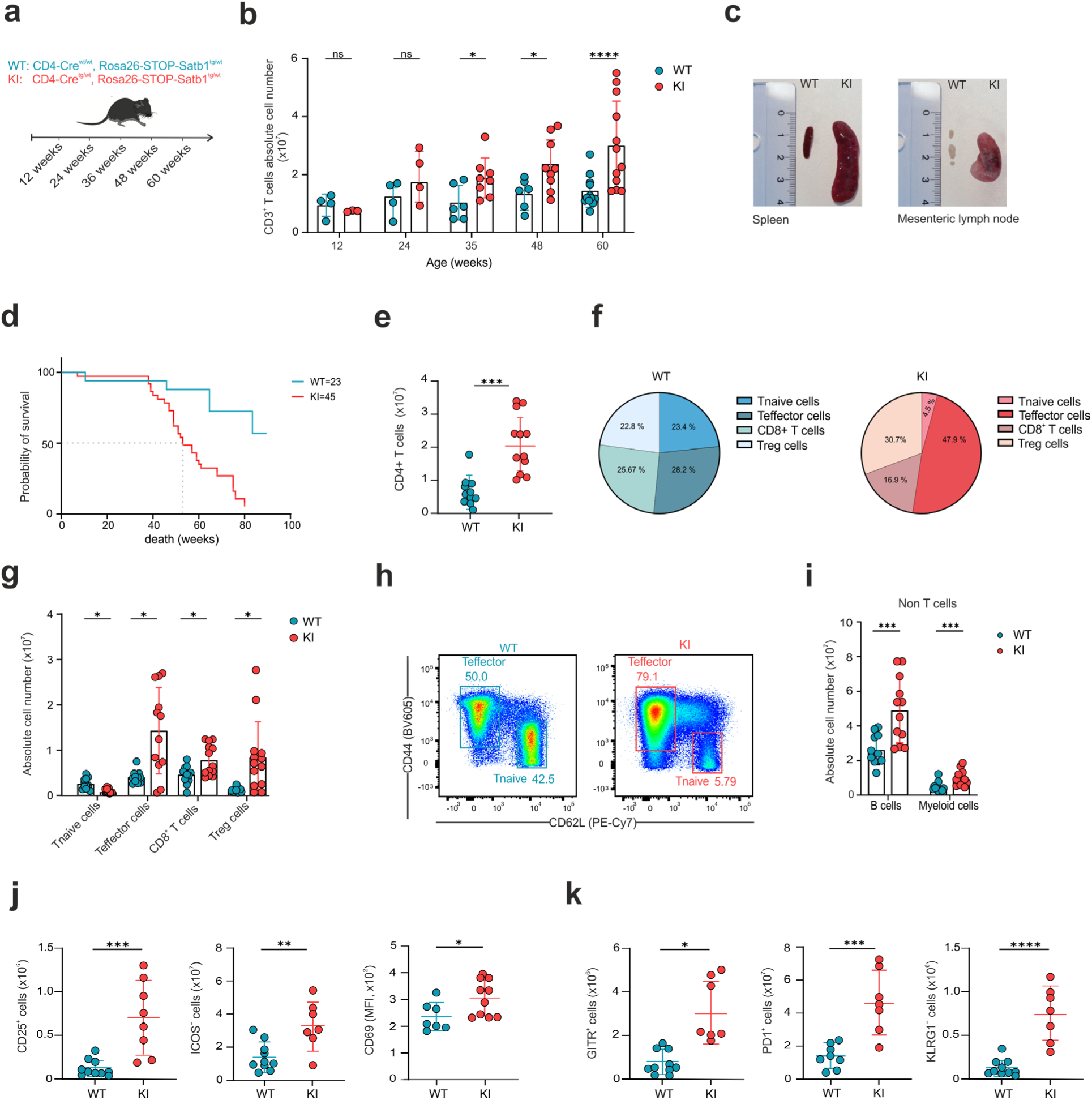
Increased Satb1 expression promotes lymphoproliferation and CD4^+^ T cell expansion in aged mice. **a,** Experimental workflow for assessing CD3^+^ T cell populations in mice sequential time points during their lifespan. **b,** Absolute cell number of splenic CD3^+^ T cell populations in mice sequential time points during their lifespan (n varies by time point). **c,** Gross morphology of spleens and mLN from the WT and KI mice, with ruler included as scale reference. **d,** Survival of WT and Satb1 KI mice under steady-state conditions. Mice were monitored over time in the absence of infection or experimental challenge (WT n=23, KI n=45). Survival curves were compared using the log-rank test. **e,** Absolute cell number of splenic CD4^+^ T cell populations in aged WT and KI mice (WT n=11, KI n=12). **f,** Frequency of splenic Tnaive, Teffector, CD8^+^ T cells and Treg cells in aged WT and KI mice. **g,** Absolute cell number of splenic Tnaive, Teffector, CD8^+^ T cells and Treg cells in aged WT and KI mice (WT n=11, KI n=12). **h,** Flow cytometric analysis of splenic Tnaive and effector cells from aged WT and KI mice. **i,** Absolute cell number of splenic B cells and myeloid cells from aged WT and KI mice (WT n=11, KI n=12). **j-k,** Flow cytometric analysis of activation marker expression in splenic CD4⁺FoxP3⁻ conventional T cells from aged WT and KI mice. **b-k,** Data are pooled from at least two independent experiments and are analyzed by unpaired *t*-test; ns indicates not significant *p* > 0.05, *p* < 0.05 = *; *p* < 0.01 = **; *p* < 0.001 = *** *p* < 0.0001 = ****

To characterize the impact of Satb1 overexpression on changes in T cell populations, we performed flow cytometry of T cell subsets isolated from the spleen and mLNs of aged WT and KI mice. We observed a significant expansion of CD4^+^ T cells, CD8^+^ T cells as well as CD3^+^CD4^+^FoxP3^+^regulatory T cells (Tregs), in aged Satb1 KI mice (**Fig. 1e-g** and **Supplementary Fig. 1e-g**). When we further subclassified CD4^+^ Tconv cells we could detect an expansion of CD4^+^CD25^-^CD44^high^CD62L^-^effector/memory T cells (Teffector cells) alongside a significant reduction in CD4^+^CD25^-^CD44^low/-^CD62L^+^ naïve T cells (Tnaive cells) (**Fig. 1f-h** and **Supplementary Fig. 1f-g**). Furthermore, we quantified non-T cell populations, including B cells and myeloid cells, and observed a significant increase in their absolute cell numbers. **(Fig 1i** and **Supplementary Fig. 1h)**.

To further investigate if Satb1 overexpression has any impact on activation markers of CD4^+^ FoxP3^-^ T cells, we assessed the expression of CD25, ICOS, CD69, glucocorticoid-induced TNFR-related protein (GITR), PD-1, and Killer cell lectin-like receptor G1 (KLRG1) by flow cytometry in both the spleen and mLNs. Notably, a significant upregulation of all activation markers was observed in Satb1 KI mice (**Fig. 1j** and **k** and **Supplementary Fig. 1i** and **j**). In summary, Satb1 overexpression in CD4^+^ T cells results in a pronounced lymphoproliferation and an increase in the number of T cells within both the spleen and mLNs, alongside an increased effector cell differentiation, increased activation marker expression and a concomitant reduction in mice survival.

### High levels of Satb1 do not alter Treg functionality and clonality of CD4^+^ Tconv cells

As we had observed that frequencies and absolute cell numbers of Treg cells were significantly increased in aged KI mice **(Fig. 1f** and **1g)** and that Satb1 repression is crucial for maintenance of the Treg cell phenotype and function ^22^ we asked how sustained Satb1 expression would affect Treg cell function. To investigate the influence of Satb1 overexpression on Treg survival and function, we assessed the expression of B-cell leukemia 2 (Bcl-2) and evaluated the suppression potential of these cells. Notably, we observed upregulation of Bcl2 expression in Tregs (**Fig. 2a**); however, Satb1 overexpression in aged KI mice did not impair the suppressive function of these cells. In other words, there was no significant change in the relative proliferation of conventional T cells (Tconv) suppressed by Tregs from KI mice compared to WT mice at varying Treg:Tconv ratios (**Fig. 2b** and **2c**). Furthermore, we examined the expression of Treg cell markers, including Cytotoxic T-lymphocyte-associated protein 4 (CTLA-4), Neuropilin-1, which enhances Treg-dendritic cell interactions ^23^, Helios, involved in T cell development and homeostasis and often described as proxy for thymic Treg cells ^24^, and Glucocorticoid-induced TNFR-related protein (GITR), a key marker for Treg cell development and activity ^24^. Despite slight increases in the expression of Neuropilin-1 and GITR in Satb1 KI mice (**Fig. 2d**), no alterations were detected in the expression of CTLA-4 and Helios, confirming that Treg cell functionality and identity remained mainly unchanged upon Satb1 overexpression.

**Figure 2:**
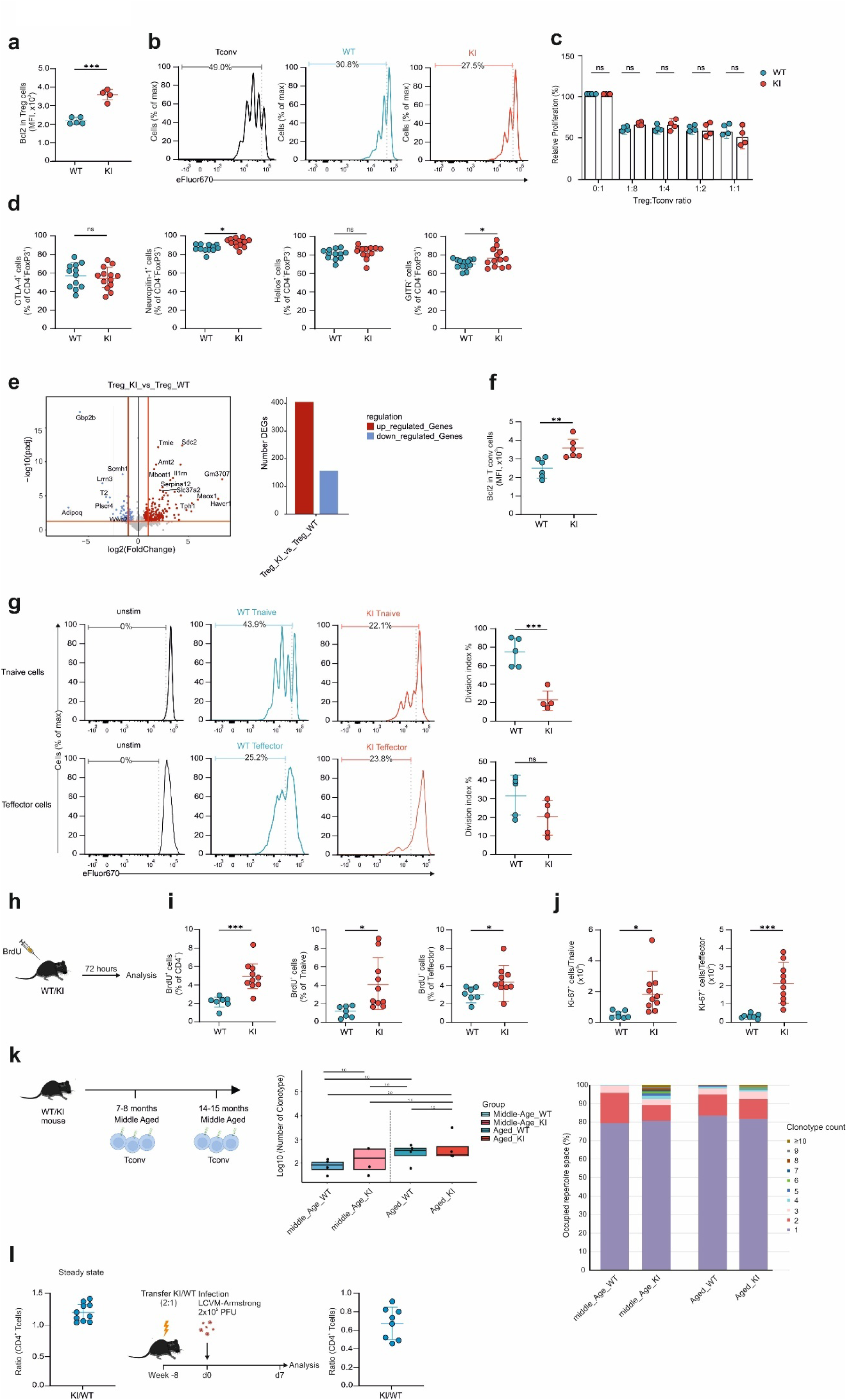
Satb1 overexpression does not disrupt T cell functionality or clonality. **a,** Intracellular Bcl2 staining of splenic CD4^+^FoxP3^+^ Treg cells from aged WT and KI mice (WT n=5, KI n=4). **b,** Suppression capacity of splenic CD4^+^FoxP3^+^ Treg cells sorted from aged WT and KI mice *in vitro*. **c,** Relative proliferation of splenic CD4^+^CD44^+^CD62L^+^FoxP3^+^ T conv cells in the presence of sorted Treg cells from aged WT and KI mice at different Treg:Tconv ratio (WT n=4, KI n=4). **d,** Flow cytometric analysis of splenic Treg markers from aged WT and KI mice (WT n=12, KI n=13). **e,** Volcano plot and summary statistic of DE genes in splenic Treg cells from aged WT and KI mice. **f,** Intracellular Bcl2 staining of splenic CD4^+^CD44^+^CD62L^+^FoxP3^-^ T conv cells from aged WT and KI mice (WT n=6, KI n=6). **g,** Proliferation of sorted CD4^+^CD44^-^CD62L^+^FoxP3^-^ Tnaive and CD4^+^CD44^+^CD62L^-^FoxP3^-^ Teffector cells from the spleen of aged WT and KI cells. **h,** Experimental workflow of BrdU experiment. **i,** Frequency of BrdU^+^ cells in CD4^+^ T cells, CD4^+^CD44^-^CD62L^+^FoxP3^-^ Tnaive and CD4^+^CD44^+^CD62L^-^FoxP3^-^Teffector cells in splenic cells from aged WT and KI cells (WT n=7, KI n=10). **j,** Absolute cell number of Ki67+ cells in Tnaive and Teffector cells from aged WT and KI mice (WT n=7, KI n=10). **k,** Bulk TCR-Seq of WT and KI CD4^+^ Tconv cells in middle-aged and aged mice. Number of unique clonotypes in the input data and summary of clonotype with specific counts. **l,** KI:WT CD4^+^ ratio at steady state and in mixed bone-marrow chimeras mice. Mixed bone marrow chimeric mice containing Satb1 KI (CD45.2^+^) and WT T cells (CD45.1/CD45.2^+^) (ratio 2:1) were created and infected with LCMV-Armstrong. **d and h**, Data are pooled from at least three independent experiments. **l,** Data are pooled from two independent experiments. **a-h**, Data are analyzed by unpaired *t*-test; ns indicates not significant *p* > 0.05, *p* < 0.05 = *; *p* < 0.01 = **; *p* < 0.001 = *** *p* < 0.0001 = ****

To further investigate the impact of Satb1 overexpression on Treg cells, we performed bulk RNA-seq of isolated CD4^+^CD44^-^CD62L^-^CD25^+^FoxP3^RFP+^ Treg cells from spleen of aged KI and WT mice. This analysis revealed 560 significantly differentially expressed genes (DE) (**Fig. 2e)**. Interestingly, we observed upregulation of *Meox1* (Mesenchyme homeobox 1), a transcription factor, whose expression in Treg cells closely resembled that of FOXP3 and could be upregulated through IL-2 stimulation in humans ^25^ (**Fig. 2e)**. Gene ontology (GO) enrichment analysis and gene–concept network plots of DE genes revealed a significant enrichment of genes associated with T cells differentiation, myeloid leukocyte activation, and organelle fission (**Supplementary Fig. 2a** and **b)**. However, heatmap visualization of Treg cell signature genes ^26, 27^ revealed that Satb1 overexpression did not change the key transcriptional genes defining Treg cell identity in KI mice compared to WT animals (**Supplementary Fig. 2c)**. Taken together, this analysis clearly demonstrated that the Treg cell functional properties and expression of the main transcriptional Treg cell programs remained unchanged upon Satb1 overexpression.

To further investigate the reason for the observed T cell expansion and shift from naïve to effector T cells in Satb1 overexpressing mice, we aimed to elucidate the factors contributing to these alterations. As anticipated, Tconv cells exhibited elevated levels of Bcl-2 in line with inhibition of apoptosis (**Fig. 2f)**. Next, we assessed T cell proliferation *in vitro*. Unexpectedly, we observed a significant reduction in the division index of Tnaive cells derived from Satb1 overexpressing mice, indicating diminished proliferative capacity, with effector T cells exhibiting a similar trend in KI mice (**Fig. 2g).** To assess whether the observed proliferative defects were inherent to T cells, we conducted adoptive transfer experiments utilizing lymphopenic Rag2⁻/⁻ recipient mice. Naive and effector CD4⁺ T cells from WT and KI donors were transferred intraperitoneally, and recipient body weight and T-cell expansion were monitored (**Supplementary Fig. 2d**). Notably, adoptive transfer into Rag2⁻/⁻ hosts demonstrated that only WT naive CD4⁺ T cells exhibited robust homeostatic proliferation, characterized by elevated Ki-67 expression, a nuclear marker of proliferation, and subsequent recipient weight loss; conversely, Satb1 overexpressing T cells displayed a markedly blunted proliferative response and failed to induce the observed wasting phenotype (**Supplementary Fig. 2d**).

To further assess T cell proliferation *in vivo*, we injected mice with BrdU (**Fig. 2h)**. We observed a significantly elevated frequency of BrdU+ cells within both Tnaive and Teffector cell populations in Satb1 KI mice (**Fig. 2i)**. Concurrent staining with Ki-67 revealed substantial upregulation of Ki-67 expression in both Tnaive and T effector cells in Satb1 overexpressing mice (**Fig. 2j)**, indicating that the Satb1 KI cells were highly proliferative *in vivo*. This prompted us to ask the critical question whether the expanded pool of T cells would result from a mono- or oligoclonal T cell origin. To investigate the clonality of conventional T cells, we performed TCR-seq of sorted T cells from aged WT and KI mice (14-15 months of age) as well as younger animals (7-8 months of age) (**Fig. 2k**). This data revealed that Satb1 overexpressing mice exhibited similar T cell clonality to WT mice, with the majority of mice displaying >70% unique clones (**Fig. 2k)**. These findings suggest that Satb1 overexpression drives *in vivo* proliferation of conventional T cells, while maintaining a polyclonal cellular landscape which is dependent on the presence of other adaptive immune cells.

To investigate whether the enhanced proliferative capacity of Satb1 KI CD4^+^ cells was intrinsic under physiologic immune conditions, we generated mixed Bone marrow (BM) chimeras reconstituted with a 2:1 ratio of Satb1 KI (CD45.2^+^) to WT (CD45.1/2^+^) bone marrow cells. This donor ratio was strategically selected, based on prior observations of diminished proliferative output *in vitro* and in Rag2 KO mice, to ensure sufficient representation of Satb1 KI cells for assessment of competitive fitness (**Figure 2l**). Following 8 weeks of reconstitution and LCMV-Armstrong infection, the ratio of Satb1 KI CD4^+^ T cells decreased relative to WT, indicating that the proliferative advantage is not a T cell intrinsic feature and likely mediated by non-cell-autonomous factors present within Satb1 KI hosts.

### Sustained Satb1 expression induces transcriptional hallmarks of Tfh and Tfr cells

Satb1 functions as a tissue-specific chromatin organizer in T cells, controlling gene expression by binding to promoter-enhancer regions and orchestrating long-range chromatin interactions, including those regulating *Bcl6* ^28^. Since Satb1 deletion reduces chromatin accessibility and disrupts the transcriptional programs critical for T cell differentiation ^29^, we hypothesized that overexpression of Satb1 enhances gene expression in naive and effector T cells. Therefore, to investigate the transcriptional programs regulated by Satb1, we performed bulk RNA-seq of sorted Teffector and Tnaive cells from WT and KI mice (**Fig. 3a**). Principal component analysis (PCA) demonstrated a clear separation between Satb1 overexpressing Teffector cells compared to WT cells while Tnaive cells were more similar, highlighting distinct transcriptional signatures driven by Satb1 (**Fig. 3b**). While SATB1 overexpression altered gene expression in both Teffector and Tnaive subsets, Teffector cells exhibited a significantly higher number of differentially expressed genes (DE genes; 1249 up, 817 down) compared to Tnaive cells (112 up, 16 down), indicating a more substantial reliance on SATB1-downregulation to enable Teffector-defining transcriptional make-up (**Fig. 3c**). Analysis of DE genes revealed that Satb1 overexpression enhances the expression of lipid metabolism genes, including *Apoe, Apoc2* and *Apoc4* (**Fig. 3d**). Further analysis revealed that SATB1 overexpression induced a distinct transcriptional program in both Tnaive and Teffector cells. Tnaive cells were characterized by the upregulation of key effector genes, including *CXCR5* while Teffector cells showed additional upregulation of *Pdcd1*, *Bcl6*, *Il2ra*, and *Ascl2*, which promote CXCR5 expression, alongside the downregulation of *Prdm1* and *Runx3* – transcriptional repressors essential for Tfh cell development ^30^ – indicating a skewing towards the Tfh cell lineage (**Fig. 3e**). Reanalysis of the Treg cells also showed similar trends supporting an increased differentiation towards Tfr cells (**Supplementary Fig. 3a**). In addition, Satb1 overexpression resulted in increased Bcl2 expression in Teffector cells, as confirmed by flow cytometry (**Fig. 3f**) with Tnaive cells showing a very similar trend (**Supplementary Fig. 3b**). To identify transcriptional programs associated with the Satb1 KI phenotype, we performed Gene set enrichment analysis (GSEA) using the human C7 immunologic signature collection ^31^, which demonstrate significant enrichment of previously reported Tfh signatures in Satb1 KI cells, indicating that Satb1KI cells adopt a transcriptional profile consistent with enhanced activation and features of early Tfh-biased or memory-like programs (**Supplementary Fig. 3c**).

**Figure 3:**
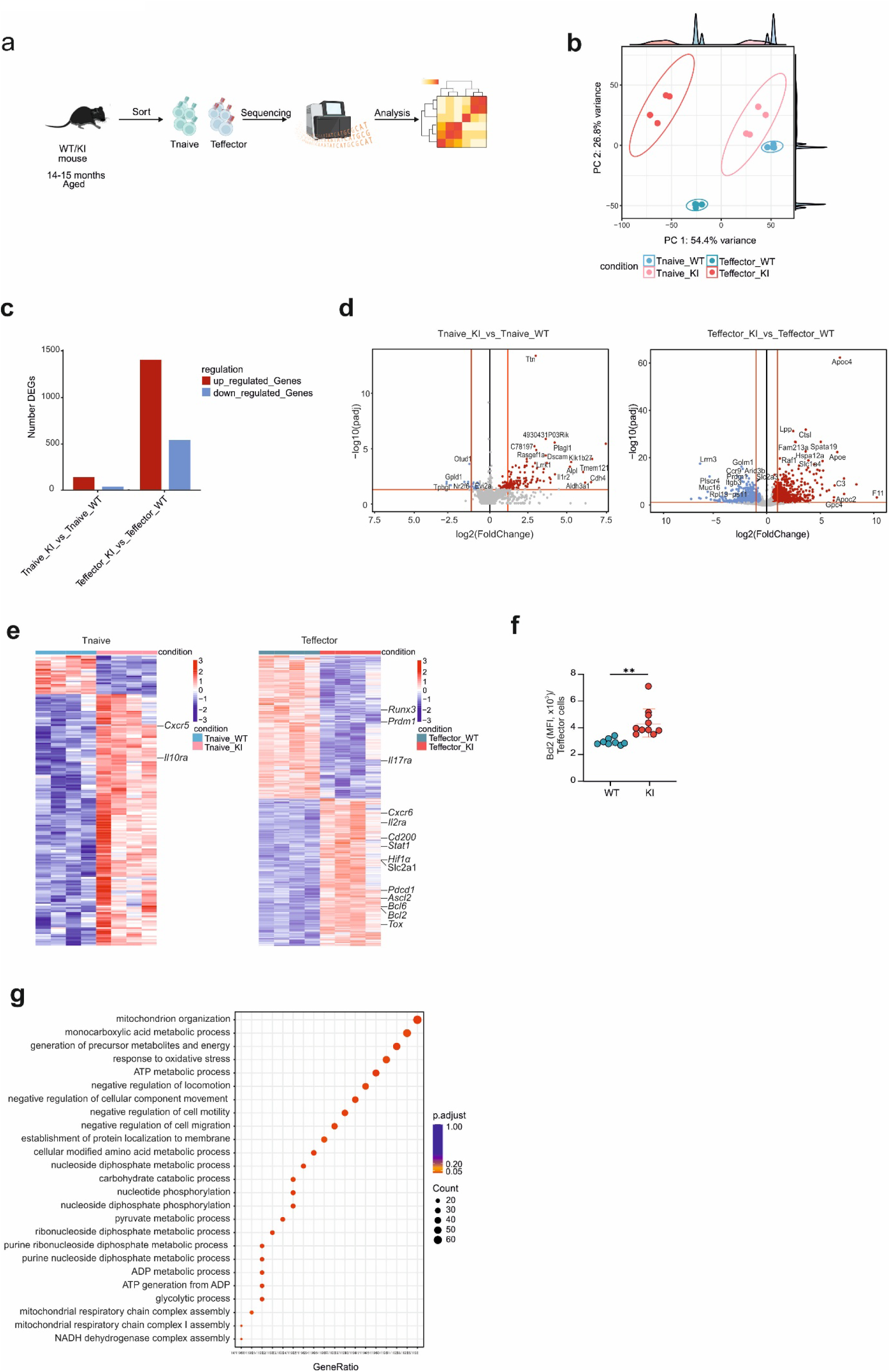
Satb1 overexpression promotes gene signatures of Tfh and Tfr cell populations. **a,** Experimental workflow for bulk RNA-seq. **b,** PCA plot of Tnaive and Teffector, cells based on gene expression data. Axes represent the first two principal components, explaining 54.4 % and 26.8 % of the variance. **c and d,** Summary statistic and volcano plot of DE genes in splenic Tnaive and Teffector cells from aged WT and KI mice **e,** Heat map showing the expression of DE genes from Tnaive and Teffector cells. Color scale indicates relative expression levels. **f,** Bcl2 expression in splenic Teffector cells from aged WT and KI mice (WT n=8, KI n=10). **g,** GO term enrichment analysis of DE genes. The size of the dots represents the number of genes in each GO category, and the color indicates the significance level (adjusted p-value).

Next, Gene ontology (GO) enrichment analysis of differentially expressed genes revealed significant enrichment of metabolic pathways, including mitochondrion organization, ATP metabolic process, and glycolytic process, alongside negative regulation of cell mobility and migration, suggesting that SATB1 influences key biological processes relevant to T cell metabolism (**Fig. 3g**). This was accompanied by increased expression of metabolic genes, notably *Slc2a1* (Glut1) and *Hif1α* upon Satb1 overexpression in Teffector cells (**Fig. 3e**) in line with previous reports demonstrating that Satb1 is indispensable for proper mitochondrial function^32, 30^.

Taken together, these data support a shift towards Tfh differentiation in CD4^+^ T cells from SATB1 KI mice and an increased metabolic activity supporting their higher proliferation *in vivo*.

### Satb1 promotes glycolytic and mitochondrial reprogramming in Tnaive and Teffector cells

Activated and quiescent T cells exhibit distinct metabolic profiles. Activated effector T cells demonstrate an anabolic metabolism characterized by high nutrient uptake and biomass synthesis, driven by glycolysis and growth factor cytokines. Conversely, quiescent T cells (naive and memory) utilize a catabolic metabolism via the TCA cycle to generate ATP from glucose, fatty acids, and amino acids and they are primarily depend on oxidative phosphorylation (OXPHOS) ^33^. To assess the functional relevance of the transcriptional enrichment of metabolic pathways observed in Satb1 overexpressing T cells, we conducted extracellular flux (XF) analysis (**Fig. 4a**). Satb1 overexpressing Tnaive and Teffector cells exhibited significantly elevated extracellular acidification rate (ECAR) and oxygen consumption rate (OCR), indicating enhanced mitochondrial respiration and glycolytic activity (**Fig. 4b, c**). These metabolic changes are consistent with the transcriptional upregulation of key metabolic regulators, including *Glut1*, *Hif1α*, *Apoe* and *Apoc4*, identified through our RNA-seq data analysis. To further validate the metabolic rewiring of SATB1 overexpressing T cells at the single-cell level, we performed single-cell energetic metabolism analysis using translation inhibition (SCENITH). This technique, which measures protein synthesis via puromycin incorporation into nascent proteins, reflects the metabolic activity of a given cell ^34^. Consistent with the XF data, Satb1 overexpressing Tnaive and Teffector exhibited significantly increased mitochondrial dependence (**Fig. 4d**).

**Figure 4:**
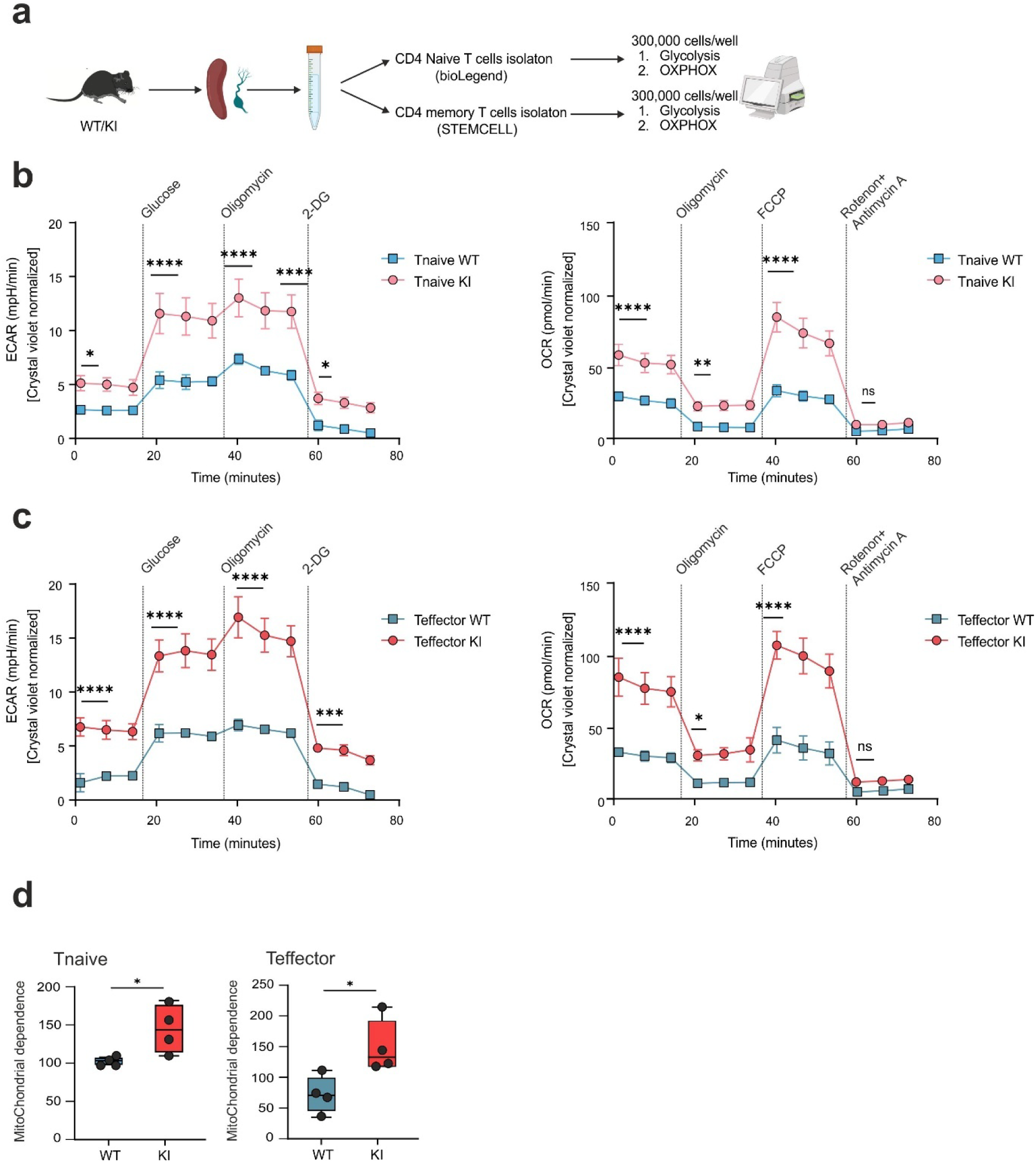
Satb1 overexpression drives enhanced glycolytic and mitochondrial activity in Tnaive and Teffector cells. **a,** Experimental workflow for Seahorse assay. **b,** Oxygen consumption rate (OCR) and Extracellular acidification rate (ECAR) of Tnaive cells from aged WT and KI mice. Data are from n = 3 mice per genotype, with technical triplicates per mouse. **c,** Oxygen consumption rate (OCR) and Extracellular acidification rate (ECAR) of Teffector cells from aged WT and KI mice. Data are from n = 3 mice per genotype, with technical triplicates per mouse. **d,** SCENITH analysis of metabolic dependencies in splenic cells from aged WT and KI cells. Data are from n = 4 mice per genotype, with technical triplicates per mouse. **b and c**, Data are analyzed by two-way ANOVA with Sidak’s test and **d,** unpaired *t*-test; ns indicates not significant *p* > 0.05, *p* < 0.05 = *; *p* < 0.01 = **; *p* < 0.001 = *** *p* < 0.0001 = ****

To further assess mitochondrial function, we utilized MitoTracker Red to measure mitochondrial membrane potential as a readout of mitochondrial activity, and MitoSOX to evaluate mitochondrial superoxide production (ROS) in Tnaive and Teffector cells from KI mice. Notably, we observed no significant difference in the frequency of MitoTracker Red and MitoSOX positive cells in either Tnaive or Teffector cell populations, indicating preserved mitochondrial function (**Supplementary Fig. 4a and 4b**). Our data strongly suggest that Satb1 overexpression in Tfh cells induces a metabolic signature encompassing both OXPHOS and glycolysis without changing mitochondrial function.

### Satb1 overexpression promotes robust expansion of T follicular helper (Tfh) and T follicular regulatory (Tfr) cells

Building upon our earlier observations of the upregulation of a Tfh and Tfr transcriptional signature, we investigated the impact of Satb1 on Th cell differentiation. Utilizing an established protocol ^35^, we differentiated Satb1 overexpressing and WT naive CD4^+^ T cells into Th0, Th1, Th2, Th17, TfhA (anti-TGF-β), Tfh, and Tfr cells *in vitro* (**Fig. 5a**). Notably, Satb1 overexpressing Tnaive cells exhibited a remarkable propensity to differentiate into Tfh cells, accompanied by a significant induction of Tfh markers (**Fig. 5b**). Expression of CXCR5, PD1, and Bcl-6 were markedly elevated in Satb1 overexpressing T cells compared to WT cells (**Fig. 5c** and **Supplementary Fig. 5a**). Furthermore, we observed a pronounced upregulation of CXCR5 and Bcl-6 in a Th17 culture condition supplemented with TGF-β and IL-6, mirroring the Tfh condition – excluding IL-21 – in Satb1 overexpressing cells. This suggests that Satb1 promotes the development of Th17 cells with enhanced Tfh-like characteristics independent of the presence of IL-21 suggesting independence of induction of Tfh cell programs from classical GC signals (**Fig. 5b**). Subsequently, we evaluated the impact of Satb1 overexpression on Th cell differentiation by analyzing *in vitro* differentiation potential of naive CD4^+^ T cells into Th1, Th2, and Th17 cells. This revealed, as previously, reported an increase in IL-17 production as well as a significant reduction in IFN-γ and IL-4 secretion (**Supplementary Fig. 5b**).

**Figure 5:**
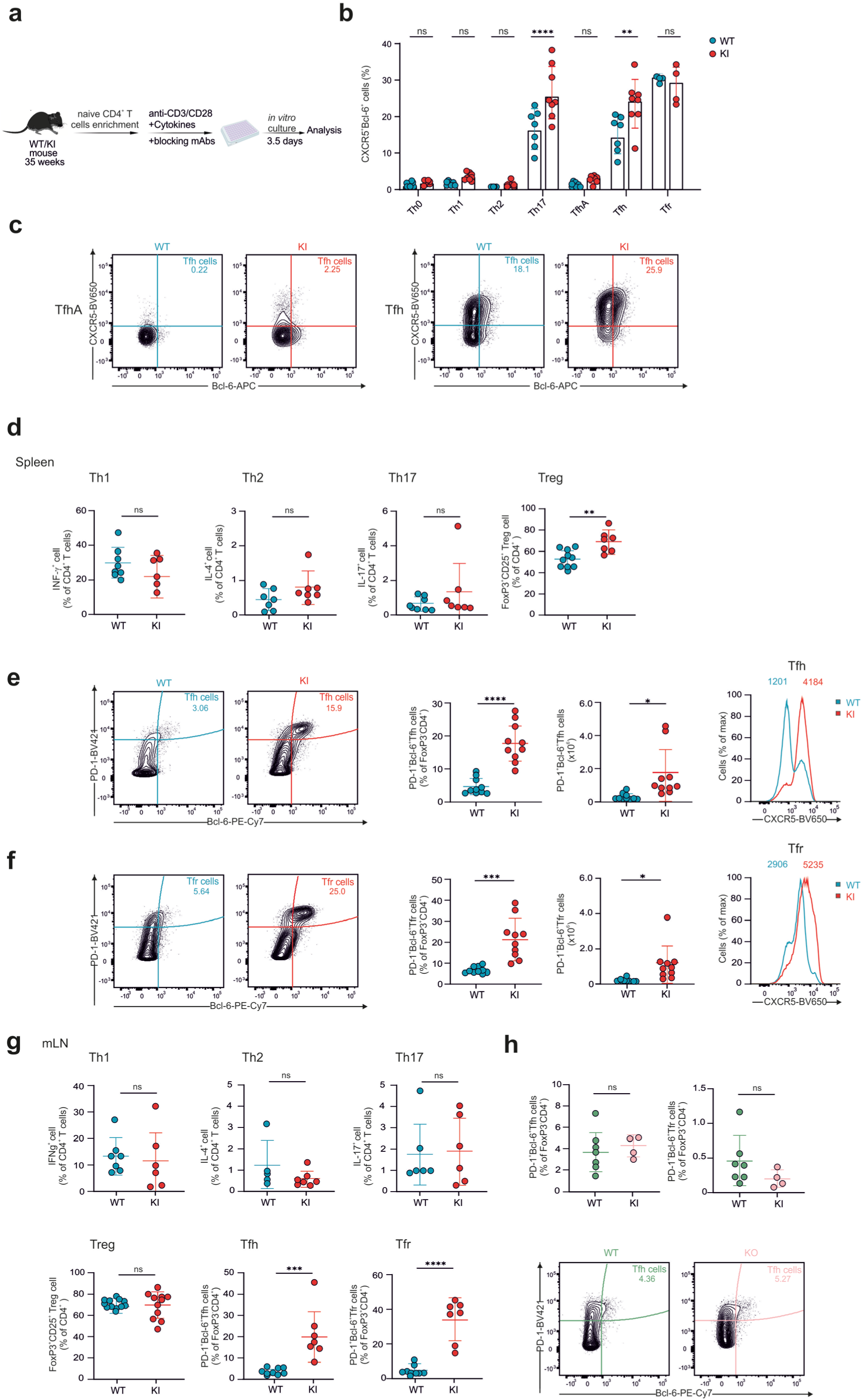
Satb1 overexpression drives expansion of Tfh and Tfr cell populations. **a,** Experimental workflow for *in vitro* Th cells subsets differentiation. **b,** Expression of CXCR5^+^Bcl6^+^ in Th subsets: Th0, Th1, Th2, Th17, TfhA (anti-TGF-β), Tfh5, Tfr cells differentiated *in vitro* from naive CD4^+^ T cells (age= 35 weeks) (WT n=7, KI n=8). **c,** Contour plot of flow cytometry showing expression of CXCR5 vs Bcl6 in TfhA and Tfh cells. **d,** *In vivo* analysis of splenic Th1, Th2, Th17, and Treg cells from aged WT and KI mice (WT n=7-10, KI n=7). **e,** Flow cytometric analysis of the percentage and total numbers of splenic Tfh cells from aged WT and KI mice (WT n=10, KI n=10). **f,** Flow cytometric analysis of the percentage and total numbers of splenic Tfr cells from aged WT and KI mice (WT n=10, KI n=10). **g,** *In vivo* analysis of Th1, Th2, Th17, and Treg cells in mLN from aged WT and KI mice (WT n=6-7, KI n=6-7). **h,** Flow cytometric analysis of the frequency of Tfh and Tfr in Satb1 deficient and Satb1 sufficient mice (WT n=7, KI n=4). **d-g,** Data are pooled from at least two independent experiments and are analyzed by unpaired *t*-test; ns indicates not significant *p* > 0.05, *p* < 0.05 = *; *p* < 0.01 = **; *p* < 0.001 = *** *p* < 0.0001 = ****

To corroborate these findings, we assessed the impact of Satb1 overexpression on CD4^+^ T cell subsets *in vivo* in aged mice. While we observed no significant alterations in the spleen in the production of IFN-γ, IL-4, and IL-17 by Th1, Th2, and Th17 cells, respectively, the frequency of FoxP3^+^CD25^+^ (Treg) cells was significantly elevated in Satb1 KI mice (**Fig. 5d)**. Consistent with our *in vitro* findings, we observed a significant augmentation in the frequency and absolute cell number of Tfh and Tfr cells in the spleen of Satb1 overexpressing mice (**Fig. 5e** and **5f)** as well as other secondary lymphoid organs such as mLN, peripheral lymph nodes (pLN), and Peyer’s Patches (PP) (**Fig. 5g** and **Supplementary Fig. 5c**).

To investigate whether Tfh and Tfr cell expansion occurred during post-thymic development, we analyzed the expression of Tfh and Tfr cell markers within the thymus and found no significant alterations in the frequency of these cells (**Supplementary Fig. 5c**). Furthermore, to determine if Satb1 overexpression also results in an aberrant accumulation of T cells in non-lymphoid organs, we assessed the frequency of Tfh and Tfr cells in the lung and observed a significant increase in both cell populations, suggesting infiltration of Tfh and Tfr cells into the lung tissue of KI mice (**Supplementary Fig. 5c**). Considering the observed expansion of CD4^+^ T cells in Satb1 KI mice, we quantified circulating cytokines in the serum of aged WT and Satb1 KI mice and observed enhanced levels of INF-γ, TNF-α, IL-2, IL-6, and IL-22 in the serum of KI mice (**Supplementary Fig. 5d**). In addition, IL-21 production was markedly increased in splenic CD4⁺ T cells from Satb1 KI mice which is the characteristic cytokine produced by follicular helper T cells (**Supplementary Fig. 5d**). This global increase suggests that sustained Satb1 expression disrupts immune homeostasis, highlighting the critical need for precise regulation of Satb1 levels to restrain inflammatory cytokine production. This was confirmed when we assessed Satb1 expression levels within Tfh cells in KI and WT mice using *ex vivo* staining. We observed that WT Tfh cells lost Satb1 expression, whereas Satb1 expression remained consistent with that observed in Tnaive cells within WT or KI mice (**Supplementary Fig. 5e**).

Based on our findings we next asked the question if a loss of Satb1 would prevent Tfh cell differentiation. To determine the role of Satb1 in this process, we examined the Tfh cell population in aged Satb1^fl/fl^ x CD4-Cre^+/wt^ mice. We observed no substantial differences in the frequency or absolute cell number of Tfh cells or Tfr between Satb1-deficient and Satb1-sufficient mice (Satb1^fl/fl^ x CD4-Cre^+/+^) (**Fig. 5h**). These findings, in contrast to previous reports^13^, demonstrate that Satb1 overexpression promotes the differentiation of both Tfh and Tfr cells, both *in vitro* and *in vivo*, while notably, Satb1 deletion did not significantly alter the populations of these cells under steady-state conditions.

### Sustained Satb1 expression reduces the frequency of antigen-specific Tfh cells

Tfh cells are essential for mounting effective humoral immune responses during viral infections, driving antibody production during acute infections and contributing to immune regulation and viral persistence in chronic infections ^36^. To assess the impact of Satb1 on Tfh-mediated antiviral immunity across effector and memory phases, 35 weeks old Satb1 KI and WT mice were analyzed following acute LCMV infection at the peak of the response (day 7) and during the memory phase (day 30) (**Fig. 6a)**. Notably, at the peak of the response, we observed comparable percentages and numbers of Tfh cells in the spleen of WT and Satb1 KI mice (**Fig. 6b)**. Furthermore, we observed a significantly reduced frequency of GP66^+^-specific Tfh cells in Satb1 overexpressing mice. Given the higher number of Tfh cells in Satb1 KI mice under homeostatic conditions, this suggests that mounting of an effective LCMV response was impaired in Satb1 KI mice. This was also reflected in the proliferative capacity of the cells as the increased Ki-67 expression in Satb1 KI mice was not maintained after LCMV infection in Tfh cells. At the memory time point (day 30), as expected, the frequency of WT Tfh cells contracted. In contrast, Satb1 KI mice maintained relatively high Tfh cell frequency although this increase did not achieve statistical significance (**Fig. 6c)**. Furthermore, no significant difference was observed in the frequency of GP66^+^-specific Tfh cells at day 30. However, proliferation capacity of Satb1 overexpressing cells significantly increased following viral clearance. Taken together, these findings suggest that Satb1 overexpression impairs generation or maintenance of LCMV-specific Tfh responses following viral challenge, thereby compromising effective antiviral immunity.

**Figure 6:**
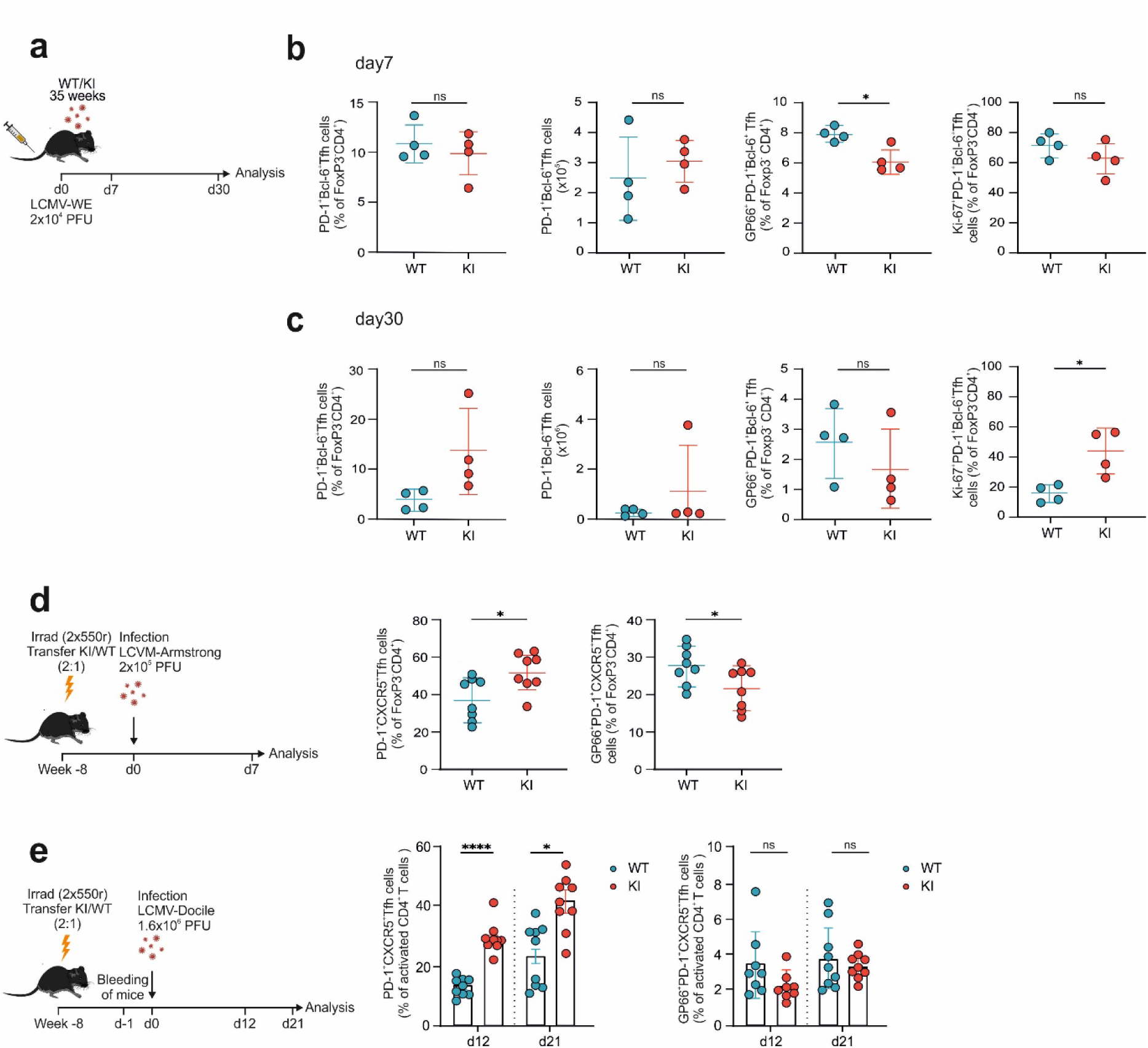
Sustained high level of Satb1 impairs the proportion of antigen-specific Tfh cells at the peak of infection. **a**, Experimental workflow for acute LCMV infection in 35 weeks old WT and mice. **b,** Flow cytometry analysis of the frequency of Tfh cells, antigen specific Tfh cells (GP66^+^), and Ki67^+^ cells on day 7 p.i. in splenic cells from WT and KI mice (WT n=4, KI n=4). **c,** Flow cytometry analysis of the frequency of Tfh cells, antigen specific Tfh cells (GP66^+^), and Ki67^+^ cells on day 30 p.i. in splenic cells from WT and KI mice (WT n=4, KI n=4). **d,** Experimental workflow and flow cytometry analysis of the frequency of Tfh cells and antigen specific Tfh cells (GP66^+^) in mixed bone marrow chimera recipient mice followed by acute infection on d7 p.i. (WT n=8, KI n=8). **e,** Experimental workflow and flow cytometry analysis of the frequency of Tfh cells and antigen specific Tfh cells (GP66^+^) in mixed bone marrow chimera recipient mice followed by chronic infection on d12 and d21 p.i. (WT n=9, KI n=9) **d, and e,** Data are pooled from two independent experiments. **a-e,** Data are analyzed by unpaired *t*-test; ns indicates not significant *p* > 0.05, *p* < 0.05 = *; *p* < 0.01 = **; *p* < 0.001 = *** *p* < 0.0001 = ****

To determine whether Satb1 dysregulation alters Tfh immunity in a cell-intrinsic manner in a fully donor-reconstituted system, we generated mixed bone-marrow chimeras by lethally irradiating recipient mice to eliminate endogenous hematopoiesis and reconstituting them with WT and Satb1 KI bone marrow at a 1:2 ratio followed by LCMV-Armstrong infection (**Fig 6d**). This approach ensured that all emerging immune cells developed in the same environment, allowing direct competitive assessment of WT versus Satb1-overexpressing T cells. At day 7 post LCMV, percentages of Satb1 KI Tfh cells reflected the transferred ratio of cells but showed reduced representation within the GP66⁺ Tfh population compared to WT cells within the same host. These findings indicate a cell-intrinsic impairment in antigen-specific Tfh priming. Having assessed the response to acute infection, we next examined how Satb1 KI and WT cells behaved during chronic infection in the same experimental setting as chronic infection imposes sustained antigenic and inflammatory pressure distinct from acute infection (**Fig 6e**). Following LCMV-Docile infection, Satb1 KI T cells exhibited markedly increased Tfh expansion relative to WT-derived cells on d12 and 21 post infection (p.i). However, despite significant increase in Tfh expansion, the frequency of Satb1 overexpressing antigen-specific GP66⁺ Tfh cells were limited and KI cells displayed a trend toward lower proliferation, on both d12 and d21 p.i., indicating impaired differentiation into functional effector Tfh cells. Collectively, these findings demonstrate that physiological Satb1 dosage is required in a Tfh cell-intrinsic manner to support productive virus-specific Tfh immunity.

### Satb1 overexpression impairs antigen-specific class-switched B cells post-immunization

As noted above, expansion of immune cells in Satb1 KI mice was not restricted to CD4^+^ T cells. Notably, significant expansion was observed in CD8^+^ T cells, B cells, and myeloid cells (**Fig. 1g** and **1i**), and confirmed by BrdU labeling, which demonstrated increased proliferation of these cell populations (**Fig. 7a**).

**Figure 7:**
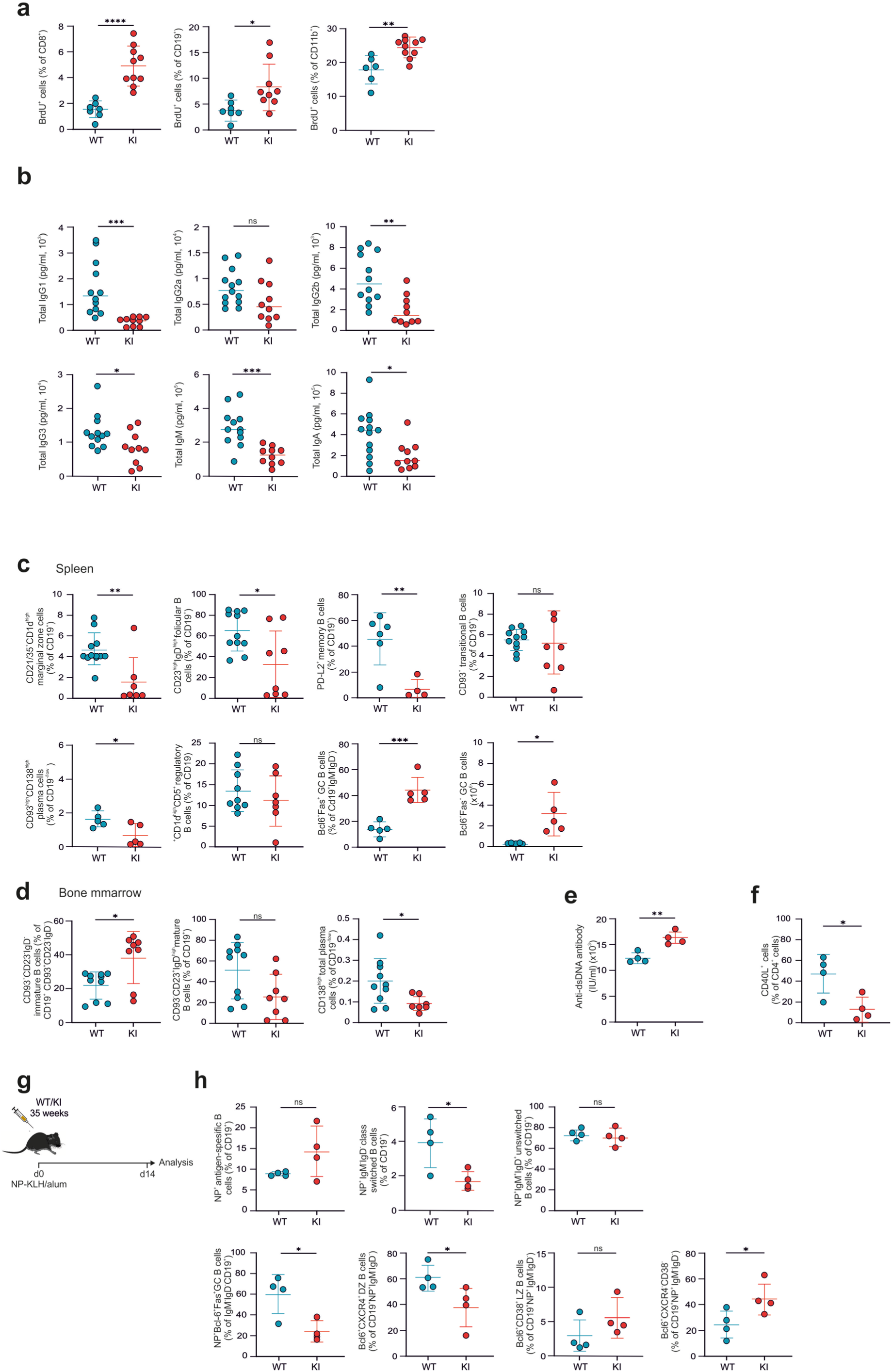
High level of Satb1 diminishes expansion of antigen-specific class-switched B cells post-immunization. **a,** Frequency of BrdU^+^ cells among splenic CD8^+^, CD19^+^, and CD11b^+^ cells from aged WT and KI cells (WT n=6-7, KI n=10). **b,** Absolute concentrations of immunoglobulin isotypes in the serum from aged WT and KI mice. **c,** Flow cytometric analysis of marginal zone cells (CD19^+^CD21/35^+^CD1d^high^), follicular B cells (CD19^+^CD23^high^IgD^high^), memory B cells (CD19^+^PD-L2^+^), transitional B cells (CD19^+^CD93^+^), plasma cells (CD19^-/low^ CD93^high^CD138^high^), regulatory B cells (CD19+CD1d^high^CD5^+^), and GC B cells (CD19^+^IgM^-^IgD^-^Bcl6^+^FAS^+^) in splenic cells from aged WT and KI cells (WT n=11, KI n=7). **d,** Flow cytometric analysis of immature B cells (CD19^+^ CD93^+^CD23^-^IgD^-^), mature B cells (CD19^+^CD93^-^CD23^+^IgD^high^) and total plasma cells (of CD19^-/low^CD138^high^) from bone marrow of aged WT and KI cells (WT n=9, KI n=8). **e,** Quantification of circulating anti–dsDNA antibodies (U/ml) in the serum of aged WT and KI mice. Data are from n = 4 mice per genotype, with technical triplicates per mouse. **f**, Flow cytometry analysis of CD40L expression in splenic CD4^+^ T cells from aged WT and KI mice (WT n=5, KI n=4). **g,** Experimental workflow for immunizing mice with NP-KLH/alum. **h,** Flow cytometric analysis of antigen-specific B cells (of CD19^+^NP^+^), class switched B cells (CD19^+^NP^+^ IgM^-^IgD^-^), unswitched B cells (CD19^+^NP^+^IgM^+^IgD^+^), GC B cells (CD19^+^NP^+^IgM^-^IgD^-^Bcl6^+^Fas^+^), DZ B cells (CD19^+^NP^+^IgM^-^IgD^-^Bcl6^+^CXCR4^+^), LZ B cells (CD19^+^NP^+^IgM^-^IgD^-^Bcl6^+^CD38^+^) in splenic cells from immunized WT and KI mice. **a, c, and d,** Data are pooled from two independent experiments. **a-h,** Data are analyzed by unpaired *t*-test; ns indicates not significant *p* > 0.05, *p* < 0.05 = *; *p* < 0.01 = **; *p* < 0.001 = *** *p* < 0.0001 = ****

Besides Tfh and Tfr cells, the GC is populated with B cells which play a central role in the adaptive immune responses. Tfh cells interact with a small proportion of B cells in the light zone (LZ) and the cells with high affinity for antigen will be selected for migration into the dark zone (DZ) to undergo somatic hypermutation (SHM) and proliferation ^37^. Given the observed expansion of B cells in KI mice, we investigated whether serum immunoglobulin levels were elevated. To address this question, we quantified serum immunoglobulin levels in WT and KI mice. Notably, we observed a significant reduction in the secretion of IgG1, IgG2b, IgG3, IgM, and IgA under steady-state conditions (**Fig. 7b**). To further characterize the expanded B cell population, we performed flow cytometry and observed a significant reduction in the frequency of marginal zone B cells, follicular B cells, memory B cells, and plasma cells, alongside a corresponding increase in GC B cells, within both the spleen and pLN of Satb1 overexpressing mice (**Fig. 7c** and **Supplementary Fig. 6a**). Analysis of the BM B cell compartment revealed an increase in immature B cells and a concomitant reduction in plasma cells (**Fig. 7d**). This shift suggests a block in the transition from immature to mature follicular or marginal zone B cells, ultimately leading to B cell inactivation and the inability to differentiate into plasma cells.

To investigate the transcriptional basis of these phenotypic alterations, we performed RNA-seq analysis on sorted B cells (**Supplementary Fig. 6b**). This analysis revealed a distinct set of differentially expressed genes in B cells from Satb1 KI mice compared to controls. Notably, upregulation of *Mki67* and *S1pr2*, genes associated with cell proliferation, was observed. Furthermore, *Aicda* (encoding AID, a key enzyme in immunoglobulin class switch recombination and SHM ^38^) was significantly upregulated in Satb1 overexpressing mice. Given that activation-induced cytidine deaminase (AID) upregulation has been reported in lupus-prone MRL/*Fas^lpr/lpr^* mice – a well-established model which develops a systemic autoimmune disease resembling human Systemic Lupus Erythematosus (SLE) ^39^ – we measured anti-dsDNA antibody secretion (key biomarkers of disease progression ^40^) and observed a surprising increase in its secretion in Satb1 overexpressing mice (**Fig. 7e**). These findings, alongside the established role of Satb1 in immune cell function, suggest that Satb1 may contribute to the development of B cell-mediated autoimmunity.

Given that reduced CD40L expression can impair T cell–dependent B cell responses ^41, 42^, we quantified CD40L expression on CD4^+^ T cells and observed significant downregulation of CD40L in Satb1 overexpressing CD4^+^ T cells, suggesting defective T cell help to B cells (**Fig. 7f**). To test whether reduced CD40L alters T cell–dependent humoral immunity and to evaluate the functional consequences of the observed transcriptional and phenotypic changes in B cells, we immunized mice with NP-KLH and tracked B cell responses during the germinal center reaction (**Fig. 7g**). We found that Satb1 overexpression resulted in a marked reduction in class-switched B cells, GC B cells, and DZ B cells, alongside an expansion of CD19^+^NP^+^IgM^-^IgD^-^Bcl6^+^CXCR4^-^CD38^-^ B cells (**Fig. 7h**). Notably, we also observed an elevated frequency of Tfh cells in Satb1 overexpressing mice, mirroring the baseline frequency (**Supplementary Fig. 6c**). These findings suggest that Satb1 overexpression despite the increased numbers of Tfh cells diminishes germinal center B cell responses and impairs class-switch recombination, potentially by driving the accumulation of abnormal B cell subsets – reflecting abortive GC reactions or defective differentiation.

Tfr cells are a type of regulatory T cells that traffic to B cell follicle and GC upon immune activation and loss of the stable suppressive anergy drives their conversion into effector-like ex-Tfr cells which cause autoimmune disorders ^43, 44^. To investigate whether Satb1 overexpression has any impact on function of Tfr cells, we assessed the expression of Helios, encoded by the *IKZF2* gene, which is essential for preserving the suppressive stability of regulatory T cells ^45^. We observed significant reduction in expression of Helios in Satb1 overexpressing Tfr cell (**Supplementary Fig. 6d**), suggesting that the Tfr cells may exhibit reduced functional activity. Previous studies have also demonstrated that 6–8 months *Ikzf2* ^−/−^ mice spontaneously developed an autoimmune phenotype with SLE-like features, including lymphocytic infiltration into non-lymphoid organs, autoantibody production, and glomerulonephritis ^45^. Taken together, the observed increase in Tfh cells, GC B cells, coupled with a reduction in the frequency of marginal zone B cells, follicular B cells, memory B cells, and plasma cells, indicates a shift towards a non functional germinal center-driven response.

## DISCUSSION

Our findings demonstrate that sustained expression of Satb1 in CD4^+^ T drives the expansion of Tfh and Tfr cells, confirmed through transcriptome analyses that revealed upregulation of the Tfh and Tfr gene signatures. High level of Satb1 expression in Tfh cells impaired generation of antigen specific Tfh cells and suggest that their functional contribution to the immune response may be limited. Furthermore, we observed an increase in GC B cells, accompanied by the suppression of marginal zone, follicular, memory, and plasma cell populations. This germinal center-driven response is associated with increased production of anti-dsDNA antibody, aligning with the established pathogenesis of SLE ^46^.

Satb1 plays a critical role in the differentiation of mature CD4^+^ T cell subsets and early B cell development. Satb1 dysregulation can lead to diverse consequences, particularly within T and B cells, highlighting its importance in maintaining immune homeostasis ^15^. Previous studies have shown that Satb1 repression enhances Tfh formation by suppressing *Icos* and impairing Tfr cell development ^13^. In contrast, our findings support that Satb1 overexpression drives the upregulation of Tfh and Tfr gene signature including *Icos*, *CXCR5*, *PD1*, *Bcl6* and the downregulation of *BLIMP1* and *Runx3* – a transcriptional repressor that inhibits CXCR5, ICOS, and CD200 ^30^. Importantly, in contrast with previous study, we did not observe any alterations in the frequency of Tfh cells in Satb1-deficient T cells at steady state. This suggests that the role of Satb1 in Tfh development may be context-dependent, potentially influenced by the specific stimulation environment – in our case, the OVA vaccination. It is possible that the inflammatory signals generated by vaccination could overrule the regulatory effects of Satb1.

Bcl-6 is a conserved multi-domain protein that functions primarily as a transcriptional repressor, shaping immune cell development and function. It is critical for germinal center formation, high-affinity antibody generation, and the regulation of Tfh and Tfr cells ^47^. Recent studies demonstrate that Satb1 directly regulates *Bcl6* expression by facilitating long-range chromatin interactions at its promoter, maintaining high transcriptional activity ^28, 48^. Disrupting these interactions in the absence of Satb1 leads to diminished *Bcl6* expression at both the RNA and protein levels, highlighting a Satb1-dependent chromatin mechanism that may prime Tfh precursor cells during early T cell development and help establish proper Tfh cell identity ^28^. This, combined with our observation that Satb1 overexpression leads to upregulation of Bcl-6, results in the expansion of Tfh, Tfr, and GC B cells.

Recent studies have begun to elucidate the role of Satb1 in GC B cells. Using a B cell–specific conditional knockout model, Thomas et. al. (2023) demonstrated that Satb1 is dispensable for early B-cell development but exerts a stage-dependent regulatory function in mature B cells, acting as an activator of immunoglobulin (Ig) gene transcription in resting B cells and a repressor in activated B cells ^49^. Within the GC compartment, Satb1 deletion resulted in enhanced SHM at Ig loci, indicating a protective role in preserving genomic integrity during affinity maturation ^49^. Complementing these findings, Ozawa et al. (2022) reported dynamic Satb1 expression during B cell maturation, with high expression in naive splenic B cells that declines upon activation ^50^. Notably, Satb1^high^ B cells within the GC exhibited features of the dark zone phenotype, and conditional Satb1 deletion led to enlarged GC follicles, suggesting a role for Satb1 in regulating GC architecture ^50^. However, unlike the findings reported by Ozawa et al. (2022), our data indicate that Satb1 overexpression results in expanded germinal center structures in spleen. In addition, we show that Satb1 overexpression increases GC B cell differentiation and immature B cell differentiation, while simultaneously reducing plasma cell expansion. Ultimately, this shift prevents B cells from completing their development and producing antibodies. Specifically, Satb1 overexpression impairs antigen-specific class-switched B cells in KI mice post-immunization, diminishing Tfh-B cell collaboration and less effective class-switch recombination, alongside impaired recruitment of B cells into the DZ for affinity maturation and increased accumulation of abnormal B cell subsets – possibly reflecting abortive GC reactions or defective differentiation.

Systemic lupus erythematosus (SLE) is a chronic autoimmune disease characterized by high levels of autoantibodies and multiorgan damage. Studies show splenomegaly develops in MRL-lpr/lpr mice, a murine model of SLE ^51, 52^. To date, there is no report examining the involvement of Satb1 in the development or progression of SLE. However, the observed increase in lymphoid organ size (mLN and spleen) and elevated secretion of anti-dsDNA antibodies, alongside AID upregulation in GC B cells in Satb1 overexpressing mice, suggests a potential role for Satb1 dysregulation in SLE development. Further studies are required to confirm this role. In agreement with the data presented here, increased glycolysis and oxidative phosphorylation in Tfh cells demonstrate that metabolic alterations, particularly increased glycolysis and oxidative phosphorylation, are implicated in lupus pathogenesis. Studies of CD4⁺ T cells from SLE patients and murine models reveal an overall heightened metabolic activity, with both pathways upregulated ^53^. In addition, studies showed Klf2-deficient mice develop spontaneous lupus-like autoimmunity by 6–8 months, with lymphocytic tissue infiltration, autoantibodies, and glomerulonephritis ^45^. In agreement with this observation, our results demonstrate that despite expansion of Tfr cells, these cells exhibited significantly diminished Helios expression, results in impaired functional activity of Tfr cells which could disrupt immune homeostasis and contribute to autoimmune pathology resembling systemic lupus erythematosus

All together, this research highlights the multifaceted role of Satb1 in immune regulation, demonstrating its critical function in both T and B cell development and a potential role in the pathogenesis of auto-antibody mediated diseases. Satb1 overexpression results in expanding Tfh and Tfr cells, GC B cell differentiation, GC expansion, enhancing autoantibody production, and splenomegaly, mirroring the pathology observed in SLE patients. Further investigation is needed to fully elucidate role of Satb1 in the pathogenesis of autoimmune diseases driven by autoantibodies, but these findings suggest a potential target for therapeutic intervention.

## Acknowledgements

We thank Michael Kraut, Heidi Theis, Dina Huesson, Svenja Bourry, and Stephanie Weber, for technical assistance.

## Author contributions

This study was conceived by M.H.S., and M.D.B. Experiments were performed by M.H.S., M.K., T.E., L.Ba. D.H., L.H., A.L., A.F., Y.I., M.G., C.K., D.M., C.O.S., R.S., X.C., A.N. Libraries were sequenced and data was pre-processed by E.D.D. Data analysis was performed by M.H.S, T.E., L.H. A.F., J. B. S., L.H. and J.S.S., Y.C., H. W., T.U., F. T. W., provided resources and critical discussions. The project was supervised by Z.A, D.B, A.K., L.B., and M.D.B. and M.H.S and M.D.B. wrote the original draft. All authors contributed to editing the manuscript.

## METHODS

### Contact for Reagent and Resource Sharing

Further information and requests for resources and reagents should be forwarded to and will be fulfilled by the Lead Contact, PD Dr. Marc Beyer.

### Mice

Mice (crossed to a C57BL/6JRcc background) were housed under specific pathogen-free conditions with chow and water provided *ad libitum.* Mice were used in accordance with the local legislation governing animal studies following The Principles of Laboratory Animal Care (NIH publication No. 85-23, revised in 1996). All animal experiments were approved by the Local Animal Care Commission of North Rhine-Westphalia or the university of Melbourne. Conditional R26-STOP-Satb1 mice were generated in house. For experiments, the following genotypes were used: wild-type CD4-Cre^wt/wt^, Rosa26-STOP-Satb1^tg/wt^, as WT and conditional CD4-Cre^tg/wt^, Rosa26-STOP-Satb1^tg/wt^ as Satb1 KI ^18^. Flow analysis utilized both male and female mice, as no significant phenotypic differences were observed. The mice were age matched. Sequencing data was generated using male mice only. For the adoptive T cells transfer, Rag2^-/-^ mice were used ^54^. The influence of long-term deletion of Satb1 in Tfh cells was studied in 60 weeks old Satb1^fl/fl^ x CD4-Cre^wt/+^ mice and Satb1^fl/fl^ x CD4-Cre^wt/wt^ as control. Mixed bone marrow chimeric mice were generated by irradiation of CD45.1^+^ recipients (2x 550 Rad). Reconstitution was done with either Satb1*^m1^*^Anu/*m1*Anu^ ^55^ or *Satb1* KI (as described before) mixed congenically marked control (CD45.1/2^+^ *Cd4*^Cre^) bone marrow cells. After reconstitution and 6-8 weeks of recovery, mice were used for further experiments.

### LCMV infection

LCMV-Armstrong and -Docile were propagated, and viral titers were quantified as previously described ^56^. Frozen stocks were diluted with phosphate-buffered saline (PBS). Each mouse was infected with 1.6×10^6^ plaque-forming units (PFU) LCMV-Docile intravenously, 2×10^4^ LCMV-WE intravenously, or 2×10^5^ PFU LCMV-Armstrong intraperitoneally.

### Immunization of Mice

35-week-old mice were immunized with 4-hydroxy-3-nitrophenol–Keyhole Limpet Hemocyanin (NP-KLH) (Biosearch Technologies, N-5060-25). NP-KLH was prepared by premixing with a 1:1 ratio of phosphate-buffered saline (PBS) and Alum (Sigma Aldrich,) prior to injection. Mice received 500 μg of the NP-KLH/Alum mixture via intraperitoneal (i.p.) injection on day 0, and were subsequently analyzed at day 14 post-immunization.

### Isolation of immune cells

Immune cells were isolated from spleen, mLN, pLN, thymus, Peyer’s patche (PP), and lung. For lymphocytes isolation from spleen, LNs, thymus, and PP, organs were meshed through a 100 μm strainer in a 50ml falcon tube. Splenocytes were spun down at 300 g 4 °C for 6 min, and supernatant was aspirated using a suction pump. In order to clear splenocytes form erythrocytes, the cells were incubated in red blood cell lysis buffer (155 mM NH4Cl, 10 mM KHCO3, 0.1 mM EDTA, pH 7.2-7.4) for 2 min. The cells were rinsed with 1x phosphate buffered saline (PBS; Sigma, D8537-500ML) and immediately used for subsequent experiments. For isolation of lymphocytes from lung, the tissue was cut into small pieces and digested in 3 mL IMDM (Gibco, Thermo Fisher Scientific, 12440053) supplemented with 10% FCS (heat-inactivated fetal calf serum), 1% non-essential amino acids (Gibco, 11140-035), 1% Glutamax (Gibco, 35050-038), 5 mM HEPES (Gibco, 15630-056), 0.5 mM sodium pyruvate (Gibco, 11360-039), 55 µM β-mercaptoethanol (Gibco, 31350-010), 100 U/ml Penicillin and 100 µg/ml Streptomycin (Gibco, 15140-122) and 50 µg/mL DNAseI (Roche, 11284932001) and 500 µg/mL Collagenase, type IV (Merck, C5138) for for 45 minutes at 37°C. Then, the cells were homogenized with 20g needle and transferred through 100µm strainer into new 50 ml falcon. The red blood cell lysis buffer was used when necessary. Cell suspension was filtered through a 40 µm cell strainer after resuspension in PBS and centrifuged at 300 g 4 °C for 6 min. Cells were used immediately for FACS staining or stimulated for determining the cytokine profile.

### Antibodies, staining and flow cytometry analysis

Fluorescent-labeled antibodies were purchased from BD Bioscience, Biolegend or Thermo Fisher Scientific. Isolated cells were transferred into a 96-well plate (Sarstedt, 83.3925500). After pelleting the cells, surface staining was performed with different combinations of fluorescently labelled antibodies targeting surface antigens including anti-mouse Fc block (BioLegend, 101319; 1:200) and fixable Live Dead viability near-infrared dye (Thermo, L34975; 1:1000) in PBS for 20 min at 4°C in the dark. For intracellular staining, cells were fixed using the eBioscience Transcription Factor Staining kit (eBioscience, 00-5523-00) according to the manufacturer instructions. Cells were resuspended in 200 µL of fixation agent for 1 hour at room temperature. After washing twice with 1x perm buffer, the cells were stained with antibodies targeting intracellular transcription factors/cytokines at room temperature for 30 minutes in the dark. The samples were acquired on BD Symphony A5. For LCMV experiments, antigen specific CD4^+^ T cells were identified using an I-A^b^ GP66–77 MHC class II tetramer. Single cells were incubated with APC-conjugated I-A^b^ GP66–77 tetramer for 2 hours at 37°C in the dark after surface staining. The cells were washed, fixed, and intracellularly stained as described before.

### Ex-vivo stimulation of T cells

Cells were stimulated with PMA/ionomycin stimulation, for 4 hr at 37 °C and 5% CO_2_ using eBioscience Cell Stimulation Cocktail (500X), and with the addition of BD Golgi Plug and Golgi Stop, according to the manufacturer’s instructions after surface staining. Cells were then washed once using PBS, followed by Live Dead viability dye (1:1000 in PBS) staining for 15 min, and fixation using the eBioscience kit and intracellular staining as described before.

### Multiplex bead-based assessment of serum antibody isotypes and cytokine profiles

Serum was collected from mice, clarified by centrifugation (10,000 × g, 5 min, RT), aliquoted, and stored at −80 °C. Pro- and anti-inflammatory cytokines were measured using the LEGENDplex Mouse Th Cytokine Panel (BioLegend, 741044) and total immunoglobulins were quantified using the LEGENDplex Mouse Immunoglobulin Isotyping Panel (BioLegend, 740493) according to the manufacturer’s protocol. Plates were acquired on flow cytometer BD Symphony A5 at low flow rate, and ≥300 events per bead population were collected. Raw FCS files were analyzed using the LEGENDplex Cloud-based Analysis Software (BioLegend).

### Histology

For histological analysis, Spleen tissue was fixed in 4% paraformaldehyde for 48 hours at 4°C and dehydrated through a graded ethanol series, cleared in xylene, and embedded in paraffin. Sections were stained with hematoxylin and eosin and assessed for disease progression, as previously reported ^57^. In short, Tissue sections were prepared utilizing a microtome to obtain paraffin-embedded tissue sections with a thickness of 4–5 µm, which were subsequently mounted onto glass slides. The preparation involved a multi-step process: initial deparaffinization utilizing xylene (two 5-minute exchanges), followed by a graded rehydration series through ascending ethanol concentrations (100%, 95%, 70%, and 50% ethanol, each for 2–3 minutes), and final rinsing in distilled water. Following this, sections were stained with hematoxylin for a 5-minute period, rinsed in running tap water, differentiated in 1% acid alcohol, and blued using Scott’s tap water substitute (or running tap water). After rinsing in distilled water, the slides were counterstained with eosin for a 1–2-minute duration, dehydrated through ascending alcohol concentrations, cleared in xylene, and finally mounted with coverslips using a permanent mounting medium. Slides were imaged using a Zeiss microscope at appropriate magnifications.

### In vitro CD4^+^ T cells differentiation

Differentiation of naive CD4^+^ T cells was performed as previously described ^35^. In brief, naive CD4^+^ T cells were enriched from spleens and peripheral lymph nodes by negative selection using the BioLegend’s MojoSort Mouse CD4 Naïve T Cell Isolation Kit (Biolegend, 480040) according to the manufacturers’ protocol. The 96-well flat-bottom tissue culture suspension plates (Sarstedt, 83.3924.500) were coated with anti-CD3 (2 μg/ml; Tonbo Biosciences, 70-0031-U500) and anti-CD28 (2 μg/ml, Tonbo Biosciences, 70-0281/U500) in 50-μl Ca2^+^-containing phosphate-buffered saline (PBS) overnight at 4°C. Plates were washed twice, and 4 × 10^4^ naive CD4^+^ T cells were cultured in 200 μl of complete medium [RPMI +10% fetal calf serum (FCS) + 10 mM Hepes + penicillin/streptomycin (100 U/ml) + 1 mM sodium pyruvate + 1× non-essential amino acid solution + 50 μM β-mercaptoethanol] at 37°C and 5% CO2 for 3.5 days. For specific differentiation, the following cytokine and antibody mix were added to the culture: Th0 (10 µg/ml anti-IL-4 [Biolegend, 504122] and 10 µg/ml anti-IFNy, [Biolegend, 505834]), Th1 (20 ng/ml IL-12 [Peprotech, 210-12] and 10µg/ml anti-IL-4), Th2 (40 ng/ml IL-4 [Peprotech, 214-14] and 10 µg/ml anti-IFNy), Th17 (50 ng/ml IL-6 [Biolegend, 575706], 5 ng/ml human TGF-β1 [Peprotech, 100-21-10UG], 10 µg/ml anti-IL-4, and 10 µg/ml anti-IFNy), and Tfh (50 ng/ml IL-6, 5 ng/ml human TGF-β1, 25 ng/ml IL-21 [Biolegend, 574504], 10 µg/ml anti-IL-4, and 10 µg/ml anti-IFNy). In TfhA cultures, TGF-β was blocked with anti- TGF-β antibodies (BioXCell, BE0057) at 10 µg/ml.

### Sorting of B cells, Regulatory T cells and naive and effector/memory CD4^+^ T cells

Splenocytes were isolated and stained with LIVE/DEAD Fixable Near-IR, anti-CD3 (Biolegend, 100234), anti-CD4 (Biolegend, 100540), anti-CD25 (Biolegend, 102008), anti-CD62L (Biolegend, 104418), anti-CD44 (Biolegend, 103020), anti-CD19 (Biolegend, 115512), and anti-CD8 (Biolegend, 100742). Naïve CD3^+^CD4^+^CD25^-^CD44^low/-^CD62L^+^ T cells, effector/memory CD3^+^CD4^+^CD25^-^CD44^high^CD62L^-^ T cells, conventional CD3^+^CD4^+^CD25^-^FoxP3^RFP-^ T cells, regulatory CD3^+^CD4^+^CD25^+^FoxP3^RFP+^ T cells, CD3^+^CD8^+^ T cells and B CD3^-^CD19^+^ cells were sorted on a BD FACS Aria III or BD FACSymphony S6 to greater than 97% purity.

### Proliferation and suppression assay

Splenocytes and lymph node cells were pooled after isolation and CD4^+^ T cells were enriched by negative selection using the MagniSort Mouse CD4 T cell enrichment Kit (Thermo Fisher Scientific, 8804-6821-74) according to the manufacturer instructions. Cells were labeled with 5 µM cell proliferation dye eFluor670 (Thermo Fisher Scientific, 65-0840-85) for 10 min at 37°C according to the manufacturer’s protocol. Subsequently CD4^+^ T cells were surface stained with LIVE/DEAD Fixable Near-IR, anti-CD3, anti-CD4, anti-CD25, anti-CD62L and anti-CD44. Conventional CD3^+^CD4^+^CD25^-^FoxP3^RFP-^ T cells and regulatory CD3^+^CD4^+^CD25^+^FoxP3^RFP+^ T cells were sorted on a BD FACS Aria III or BD FACSymphony S6 to greater than 97% purity. For proliferation assay, Cells were stimulated for 3 days with mouse T-Activator CD3/CD28 Dynabeads (Thermo Fisher Scientific, 11456D) in a 3:1 cell/bead ratio and subsequently dilution of eFluor670 was measured by flow cytometry. For suppression assay, Treg cells were co cultured with different ratio of Tconv cells with mouse T-Activator CD3/CD28 Dynabeads in a 3:1 cell/bead ratio and subsequently dilution of eFluor670 was measured after 3 days by flow cytometry.

### Bulk RNA-Seq analysis

Total RNA was extracted from sorted Naïve CD3^+^CD4^+^CD25^-^CD44^low/-^CD62L^+^ T cells, effector/memory CD3^+^CD4^+^CD25^-^CD44^high^CD62L^-^ T cells, regulatory CD3^+^CD4^+^CD25^+^FoxP3^RFP+^ T cells, and CD3^-^CD19^+^ B cells using miRNeasy Micro Kit (Cat no. 217084) following the manufacturer’s protocol, and libraries were prepared bulk-reaction Smart-seq2 protocol. Size distribution of the libraries was checked using the Agilent high sensitivity D1000 assay on a TapeStation 4200 system (Agilent technologies). Sequencing was performed on an Illumina NovaSeq 6000 system using NovaSeq_S1 v1.5 to generate single end 75bp reads. Raw reads were quality-checked using FastQC. For samples sequenced across multiple lanes, FASTQ files corresponding to the same biological sample were first merged prior to alignment. Transcript quantification was performed using kallisto (v 0.44.0). Alignment statistics for each sample were summarized with MultiQC (v1.5) and incorporated into the sample metadata table for downstream visualization and quality assessment. Transcript-level abundances were estimated with kallisto (v 0.44.0)^58^. Estimated counts and transcript lengths were imported into R (v4.1.0) using the tximport (v1.20.0) package ^59^ and summarized to gene-level count matrices for use with DESeq2 (v1.30.1). Gene Ontology (GO) was performed using clusterProfiler (v4.0.5) on significantly up- or down-regulated genes. Data visualization, including PCA plots, volcano plots, and heatmaps, was performed with ggplot2 (v3.4.4) and pheatmap (v1.0.12) packages in R.

### Bulk TCR sequencing analysis

Splenocytes were isolated and stained with LIVE/DEAD Fixable Near-IR, anti-CD3, anti-CD4, anti-CD25, anti-CD62L and anti-CD44. Conventional CD3^+^CD4^+^CD25^-^FoxP3^RFP-^ T cells from middle age and aged mice were sorted on a BD FACS Aria III or BD FACSymphony S6 to greater than 97% purity. Bulk TCR sequencing was performed after library preparation from isolated RNA using NEBNext® Immune Sequencing Kit (Mouse) (NEB, #E6330S) according the manufacturer’s protocol. T cell receptor (TCR) repertoire sequencing data were analyzed using the MiXCR (v4.3.0) and nf-core/airrflow (v4.3.0) pipelines. Raw sequencing reads were processed using AIRRflow, a Nextflow-based workflow configured with default parameters (https://nf-co.re/airrflow/4.3.0/), for quality control and adapter trimming. MiXCR was then utilized for clonotype assembly, V(D)J gene segment alignment, and annotation, employing the recommended NEBNext® BCR profiling settings (https://mixcr.com/mixcr/guides/nebnext-bcr/). Clonal diversity, CDR3 length distribution, and V(D)J usage were assessed from MiXCR output, supplemented with custom R scripts for data visualization and further statistical analysis.

### Analysis of metabolic parameters

Real-time metabolic parameters were measured using a Seahorse XFe96 Analyzer (Agilent) according the manufacturer’s protocol. In Brief, CD4^+^ Tnaive and Teffector cells were isolated using the MojoSort™ Mouse CD4 Naïve T Cell Isolation Kit (Biolegend, 480040) and EasySep™ Mouse Memory CD4^+^ T Cell Isolation Kit (STEMCELL, 19767), respectively. The cells were seeded on poly-L-lysine-coated (100 *μ* g/ml, Merck, P5899) Seahorse cell culture microplates (Agilent, 101085-004) with a concentration of 300,000 cells/well. Cells were then incubated for one hour in Seahorse assay medium (Seahorse XF RPMI Medium supplemented with 2mM L-glutamine, 1mM sodium pyruvate, adjusted to pH 7.4 prior to the assay) at 37◦C in non-CO2 incubator. The extracellular acidification rate (ECAR) and oxygen consumption rate (OCR) with three measurement cycles recorded at baseline and following each sequential injection of metabolic inhibitors. Each measurement cycle was defined as three minutes mixing time and three minutes measurement time.

Energy metabolism with single-cell resolution was functionally profiled using the SCENITH as previously described ^34^. Briefly, freshly isolated splenocytes were incubated with protein synthesis inhibitor puromycin (Sigma-Aldrich, P7255-100MG) in the presence or absence of metabolic inhibitors: 2-deoxy-D-glucose (2-DG) (Sigma-Aldrich, D6134-25G) and oligomycin (Sigma-Aldrich, 75351-5MG). The cells were stained with surface markers followed by fixation (30 minutes at room temperature). The cells were washed and permeabilized by incubation in 1x Perm/Wash Buffer. Subsequently, cells were blocked and incubated with anti-puromycin monoclonal antibody (EMD Millipore, MABE343-AF647) for 3 hours at room temperature in the dark, followed by washing with 1x Perm/Wash Buffer. Finally, cells were resuspended in 80 µL FACS buffer and stored at 4°C until measurement.

### MitoTrackerR Red and MitoSOX™ staining

Splenocytes were stained with MitoTrackerR Red CMXRos (ThermoFisher, M7512) and MitoSOX™ Mitochondrial Superoxide Indicators (ThermoFisher, M36008) according to the manufacturer’s protocol. In short, cells were transferred into a 96-well plate and surface staining was performed by adding 50 μL of antibody cocktail to each well, followed by incubation for 20 minutes at 4°C in the dark. Cells were then washed once with 200 μL of Hank’s Balanced Salt Solution (HBSS, Thermo Scientific, 14025100). The cell pellet was gently resuspended in 100 μL of the pre-warmed MitoTrackerR Red or MitoSOX™ staining solution and incubated for 30 minutes at 37°C in the dark. Following incubation, cells were washed twice with pre-warmed HBSS and finally resuspended in 200 μL of HBSS for flow cytometry analysis.

### Quantification and statistical analysis

Statistical analyses, excluding sequencing data, were conducted using GraphPad Prism software v10 (GraphPad Software). For comparisons between two groups, two-tailed unpaired Student’s t-tests were employed. When analyzing three or more groups under similar conditions, one-way ANOVA with Dunnett’s or Tukey’s multiple comparison tests were utilized. Two-way ANOVA was performed to assess differences between multiple groups in the context of varying conditions. Post-hoc analyses were performed dependent on the experimental design. A p-value of less than 0.05 was considered statistically significant (ns indicates not significant *p* > 0.05, *p* < 0.05 = *; *p* < 0.01 = **; *p* < 0.001 = *** *p* < 0.0001 = ****). Descriptive statistics, the performed statistical tests as well as the number of samples are stated in the figure legends. Sequencing data were analyzed using R Studio.

