## Supplementary Figures for "Downregulation of Satb1 is required to prevent autoimmunity by maintaining Tfh homeostasis"

### Supplementary Figure 1

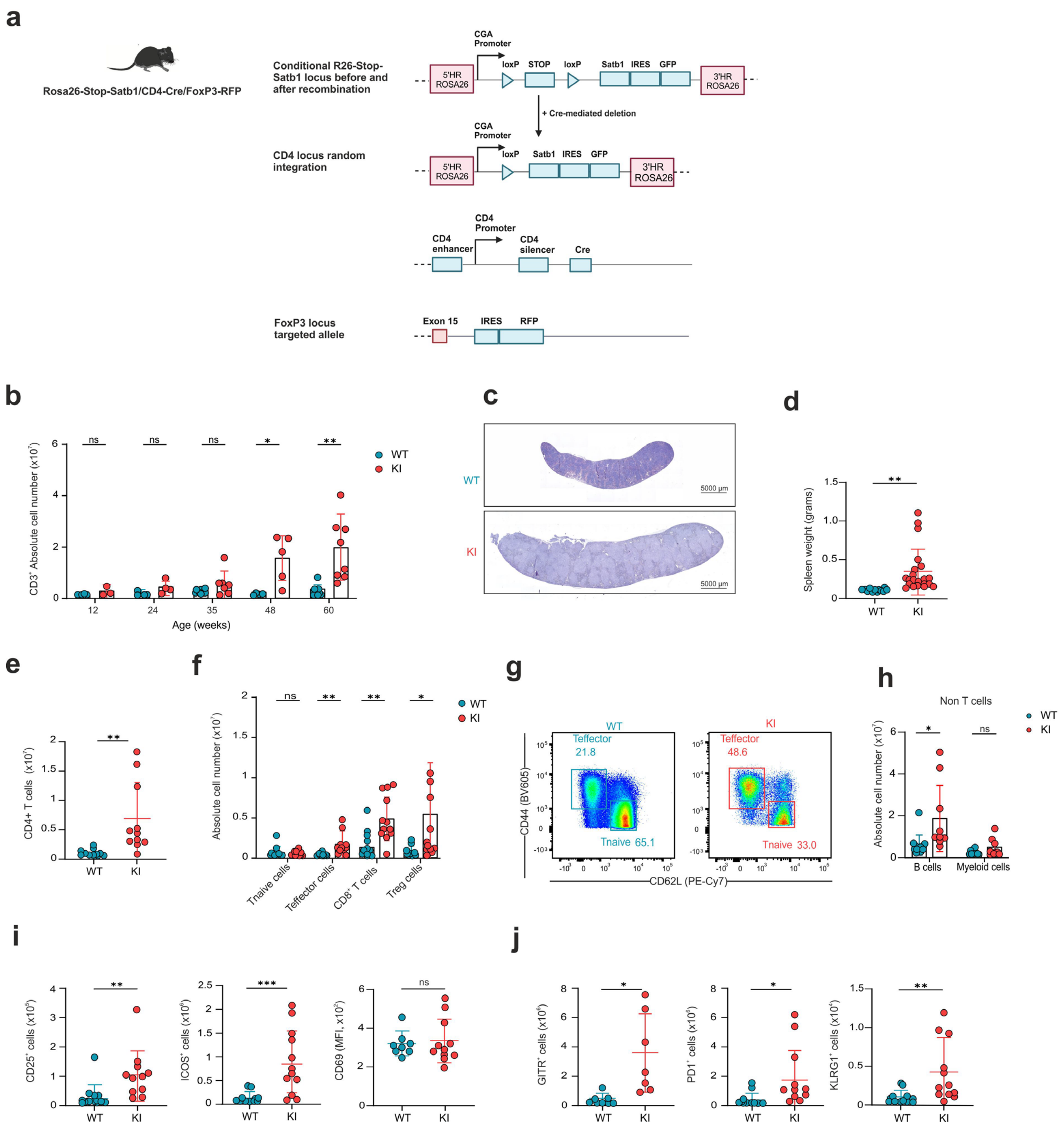

**Supplementary Figure 1: Satb1 drives CD4+ T cell expansion and lymphoproliferation in lymphoid organs of aged mice**

**a,** Schematic overview of the conditional *R26-STOP-Satb1* allele before and after Cre mediated recombination.

**b,** Absolute cell number of $CD3^+$ T cell populations in mLN across the lifespan (n varies by time point).

**c,** H&E staining sections of spleen from aged WT and KI mice.

**d,** Spleen weight of aged WT and KI mice (WT n=18, KI n=21).

**e,** Absolute cell number of $CD4^+$ T cell populations in mLN of aged WT and KI mice (WT n=10, KI n=11).

**f,** Absolute cell number of Tnaive, Teffector, $CD8^+$ T cells and Treg cells in mLN of aged WT and KI mice (WT n=10, KI n=11).

**g,** Flow cytometric analysis of Tnaive and effector cells in mLN of aged WT and KI mice.

**h,** Absolute cell number of B cells and myeloid cells in mLN of aged WT and KI mice (WT n=10, KI n=9).

**i-j,** Flow cytometric analysis of activation marker expression in $CD4^+FoxP3^-$ conventional T cells from mLN of aged WT and KI mice.

**b-j,** Data are pooled from at least two independent experiments and are analyzed by unpaired *t*-test; ns indicates not significant $p > 0.05$, $p < 0.05$ = *; $p < 0.01$ = **; $p < 0.001$ = *** $p < 0.0001$ = ****

### Supplementary Figure 2

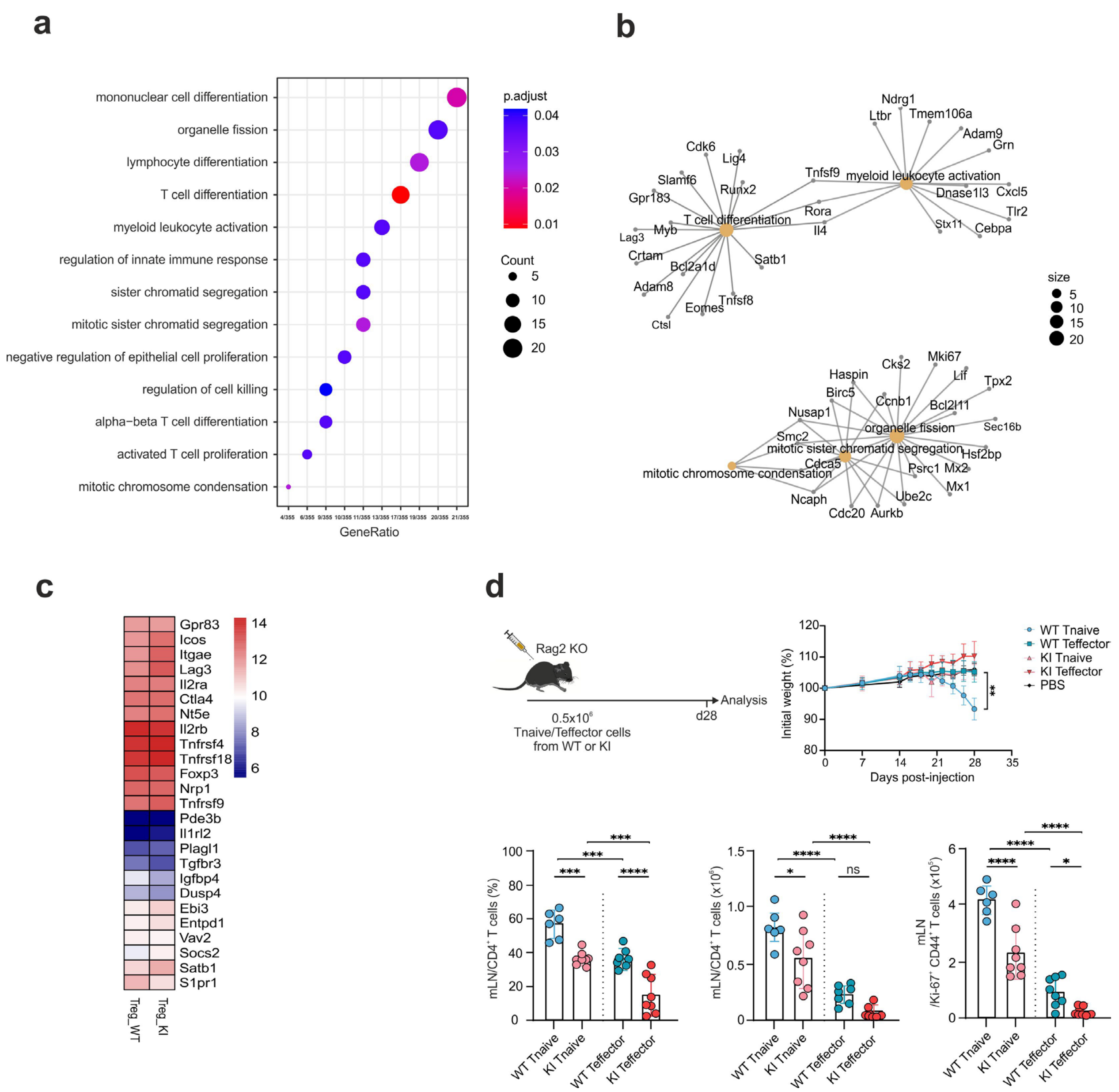

**Supplementary Figure 2: Satb1 overexpression does not change the transcriptional landscape of Treg cells**

**a,** GO term enrichment analysis of DE genes in Tregs. The size of the dots represents the number of genes in each GO category, and the color indicates the significance level (adjusted p-value).

**b**, Gene–concept network plot (cnetplot) illustrating the relationship between enriched Gene Ontology (GO) biological processes and the differentially expressed genes identified in Satb1 overexpressing Treg cells compared to WT controls. Orange nodes represent significantly enriched GO terms, gray nodes represent genes, and edges indicate the association of genes with corresponding functional categories.

**c,** Heatmap showing the mean expression of selected Treg genes from aged KI and WT mice, colored from blue to red.

**d,** Experimental workflow and proliferation of transferred WT and Satb KI Tnaive and Teffector cells from aged mice in $Rag2^{-}/^{-}$ mice (10-12 weeks) (WT n=6-7, KI n=8).

### Supplementary Figure 3

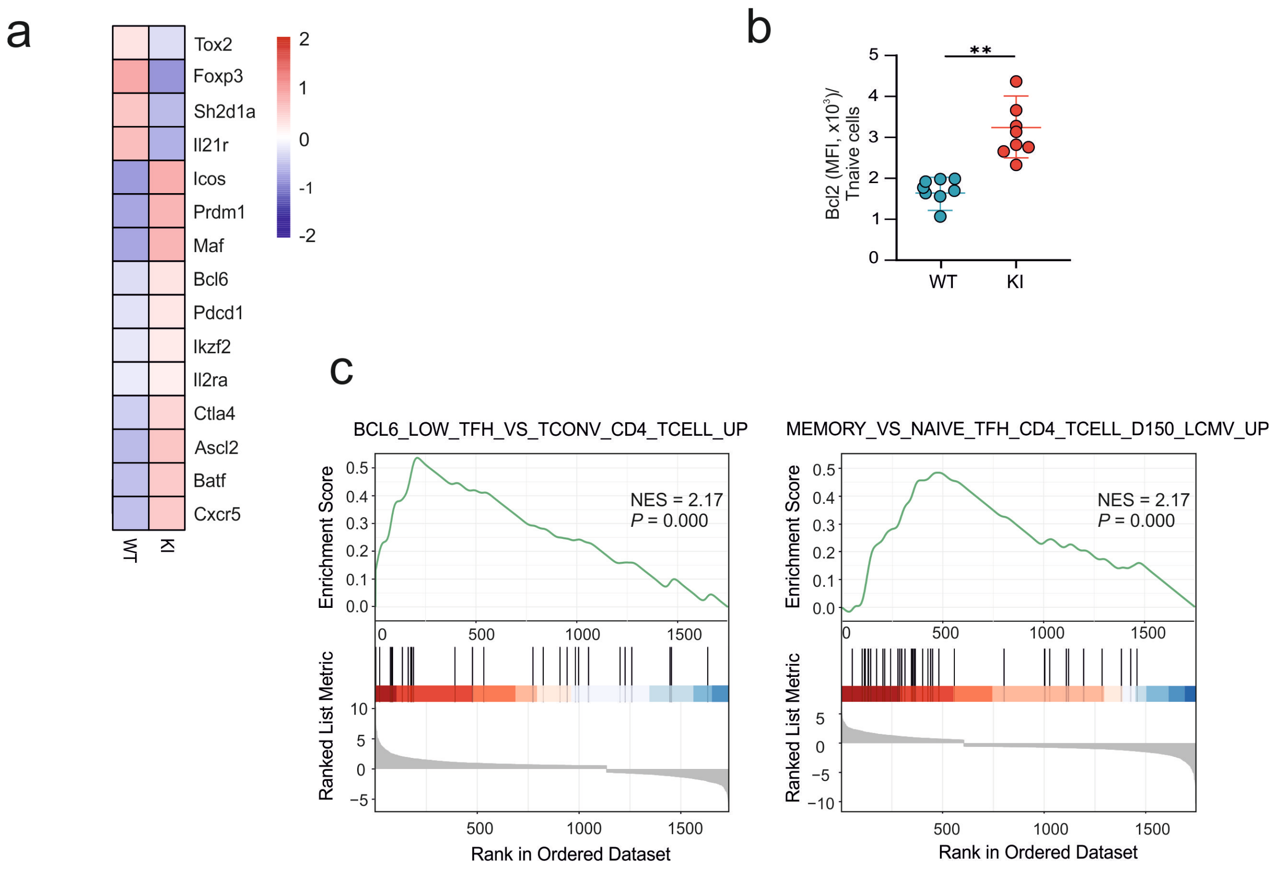


**Supplementary Figure 3: Enrichment of Tfh-associated and activation-related transcriptional programs in Satb1 KI cells**

**a,** Heatmap of Tfr signature genes showing relative expression in WT and KI samples. Gene expression was z-scored per gene across all samples and subsequently averaged within each group. Colors represent relative expression (z-score), with red indicating higher and blue indicating lower expression.

**b,** Bcl2 expression in splenic Tnaive cells from aged WT and KI mice (WT n=8, KI n=8).

**c,** GESA of the Satb1 KI versus WT cells using the human C7 immunologic signatures.

### Supplementary Figure 4

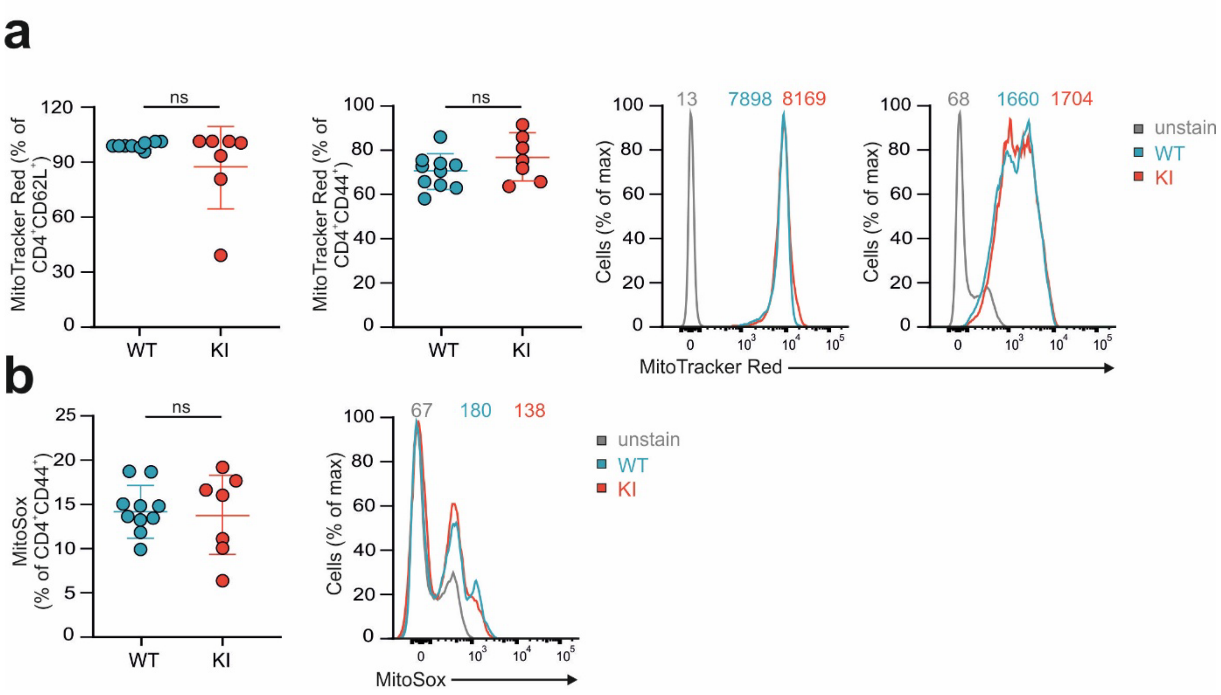


**Supplementary Figure 4: Satb1 overexpression do not alter mitochondrial function**

**a and b,** Flow cytometric analysis of MitoTracker Red and MitoSOX in splenic Tnaive and Teffector cells from aged WT and KI mice (WT n=10, KI n=7).

**a and b,** Data are analyzed by unpaired *t*-test; ns indicates not significant $p > 0.05$, $p < 0.05$ = *; $p < 0.01$ = **; $p < 0.001$ = *** $p < 0.0001$ = ****

### Supplementary Figure 5

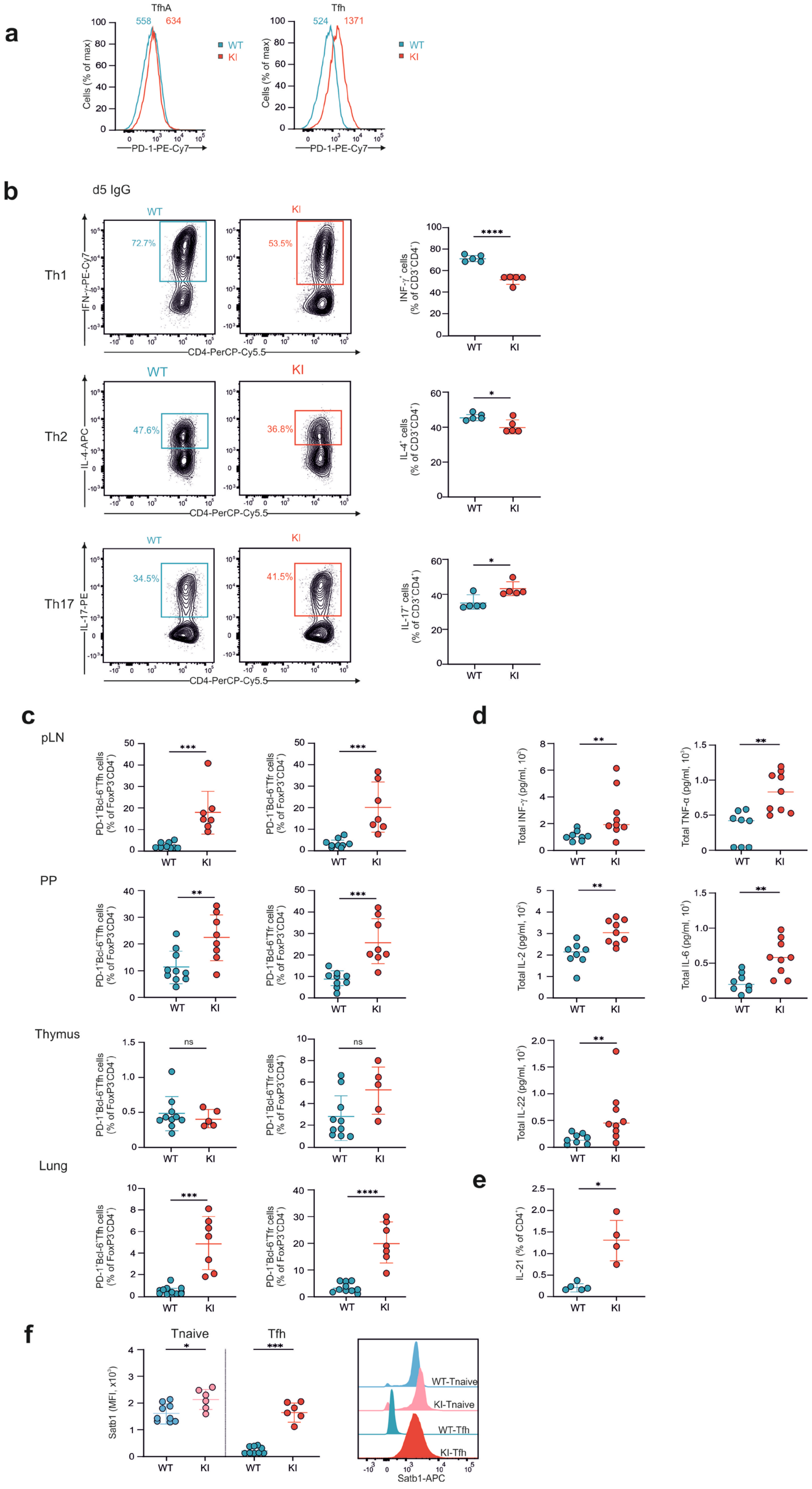

**Supplementary Figure 5: Satb1 overexpression results in Tfh and Tfr differentiation in secondary lymphoid organs**

**a,** Flow cytometry histogram plots of CXCR5 expression in TfA (anti-TGF-β) and Tfh5 cells subsets.

**b,** Analysis of *in vitro* differentiation of Tnaive cells into Th1, Th2, Th17 on day five of differentiation (WT n=5, KI n=5).

**c,** Flow cytometric analysis of the percentage and total numbers of Tfh and Tfr cells in pLN, PP, thymus, and lung from aged WT and KI mice (WT n=10, KI n=5-8).

**d,** Absolute concentrations of cytokines in the serum of aged WT and KI mice (WT n=8, KI n=9).

**e,** Flow cytometry analysis of expression in splenic $CD4^+$ T cells from aged WT and KI mice (WT n=5, KI n=4).

**f,** Flow cytometry analysis of Satb1 expression in splenic cells from aged WT and KI mice *ex vivo* (WT n=9, KI n=6).

**c** and **f,** Data are pooled from at least two independent experiments and **(b, c, and)** are analyzed by unpaired *t*-test; ns indicates not significant $p > 0.05$, $p < 0.05$ = *; $p < 0.01$ = **; $p < 0.001$ = *** $p < 0.0001$ = ****

### Supplemantary Figure 6

**a**

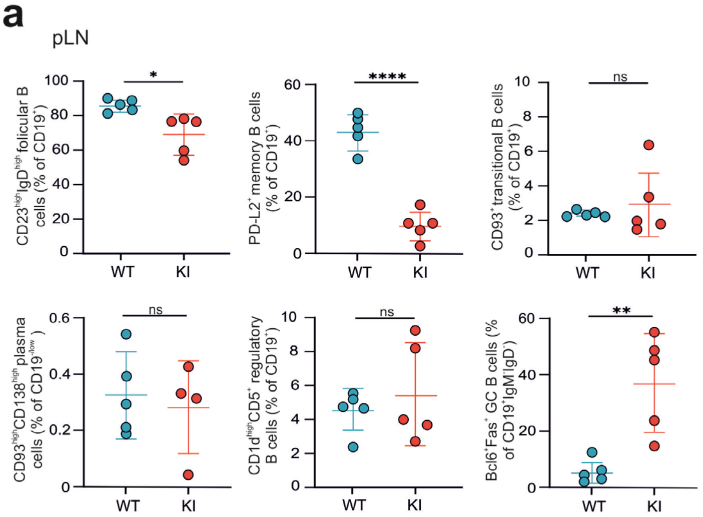


**b**

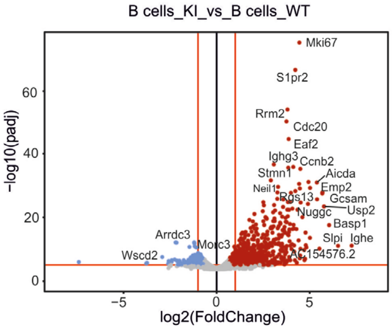


**c**

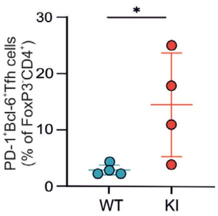


**d**

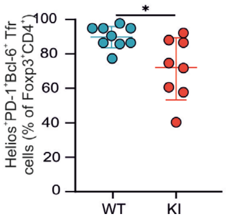

**Supplementary Figure 6: Satb1 overexpression leads GC B cells differentiation**

**a,** Flow cytometric analysis of marginal zone cells ($CD19^{+}CD21/35^{+}CD1d^{high}$), folicular B cells ($CD19^{+}CD23^{high}IgD^{high}$), memory B cells ($CD19^{+}PD\text{-}L2^{+}$), transitional B cells ($CD19^{+}CD93^{+}$), plasma cells ($CD19^{-/low}$ $CD93^{high}CD138^{high}$), regulatory B cells ($CD19+CD1d^{high}CD5^{+}$), and GC B cells ($CD19^{+}IgM^{-}IgD^{-}$ $Bcl6^{+}FAS^{+}$) in pLN of aged WT and KI cells (WT n=5, KI n=5).

**b,** Volcano plot showing DE genes in B cells from aged WT and Satb1 KI cells .

**c,** Flow cytometric analysis of splenic Tfh cells from immunized WT and KI mice. (WT n=4, KI n=4).

**d,** Flow cytometric analysis of splenic $Helios^{+}$ Tfr cells from aged WT and KI mice (WT n=9, KI n=8).

**a,b and c,** Data are analyzed by unpaired *t*-test; ns indicates not significant $p > 0.05$, $p < 0.05$ = *; $p < 0.01$ = **; $p < 0.001$ = *** $p < 0.0001$ = ****
